# Visualizing zinc-starved slow-replicating *Mycobacterium tuberculosis* in neutrophil-rich lung lesions

**DOI:** 10.64898/2026.08.21.746198

**Authors:** Jamie H. Corro, Jordan Holl, Varsha Patil, Poornima Sankar, Yunlong Li, Anas Saleh, Amy Wu, Richard W. Cole, Kyu Y. Rhee, Bibhuti B. Mishra, Anil K. Ojha

## Abstract

The months-long treatment for tuberculosis is partially attributed to a minor *Mycobacterium tuberculosis* (Mtb) subpopulation exhibiting phenotypic drug resistance, although origins of these bacilli in hosts remain unknown. Previously, we observed that mycobacteria respond to growth-restrictive limitations of zinc by accumulating Mrf, and inducing Mpy recruitment to the decoding center of the ribosome, thereby hibernating the ribosome in an inactive state. Mrf accumulation – regulated at both transcriptional and post-translational levels by Zur and Clp, respectively, in a zinc-dependent manner – facilitates the formation of stable Mpy-70S ribosome complex in zinc-starved cells. Here, using a translational fusion of Mrf with a fluorescent-reporter protein we visualized zinc-starved mycobacteria with a physiology conducive for Mpy-dependent ribosome hibernation. About half of zinc-starved mycobacterial population expressed microscopically detectable levels of the reporter and exhibited Mpy-dependent slowdown of growth and isoniazid tolerance *in vitro*. In chronic, high-burden mouse lung infections, ∼3% of Mtb bacilli expressed the reporter, and the frequency increased to ∼35% upon isoniazid treatment. Transcriptomics of mouse lungs suggests an important role of S100A8/A9 and ZIP2 in neutrophils in zinc mobilization during Mtb infections. Moreover, depletion of neutrophils, or loss of S100A9 caused significant attenuation of zinc starvation response in Mtb. Altogether, the study offers a new approach for visualizing and manipulating slow-replicating Mtb in host tissues.

**Significance:** The months-long treatment for tuberculosis is partly attributed to a minor *Mycobacterium tuberculosis* (Mtb) subpopulation exhibiting phenotypic drug tolerance. While *in vitro* models have demonstrated a correlation between drug tolerance and metabolic slowdown, origins and locations of such drug tolerant bacilli in hosts remain unknown. Here, we report a fluorescent-reporter-based method to visualize a subpopulation of Mtb bacilli in mouse lungs with slowed growth and increased tolerance to isoniazid, a drug that primarily targets growing bacilli. This approach can further lead to a better understanding of the host conditions underlying the development of these bacilli, and develop new therapeutic approaches.

## Introduction

The high global incidence of tuberculosis (TB) is partly due to months-long treatment regimens, shortening of which remains an unmet challenge (1, 2). Independent clinical trials evaluating patients’ response to TB drug regimens have identified two categories of treatment response: a) easy-to-treat (ETT) patients representing ∼80% of infections that are characterized by minimal non-cavitary disease and require less than 4 months of treatment, and b) ∼20% hard-to-treat (HTT) patients with high bacterial burden and severe lung pathological features that require longer treatment (3). This correlation between drug recalcitrance and pathological features has also been observed in experimental mouse models of TB: a C3HeBFeJ strain with extensive caseated necrotizing lesions harboring extracellular *Mycobacterium tuberculosis* (Mtb) bacilli is more refractory to drug treatment than an infected BalB/c strain with non-necrotizing lesions harboring mostly intracellular bacilli (4). Moreover, bacterial clearance in patients undergoing treatment follows a biphasic pattern (5), indicating that the extended regimen is likely necessary to eradicate a small subpopulation of bacilli that remain genetically susceptible to antibiotics (6). Such subpopulations exhibiting phenotypic resistance to antibiotics, called persisters, can be reproduced in multiple *in vitro* models, and are associated with a nonreplicating physiological state of mycobacteria (2, 7–11). Although drug resistant nonreplicating Mtb subpopulations have been documented in caseated lung lesions of infected rabbits (12) and in human sputa (13), their origin remains unclear.

Physiological adaptation in non- or slow-replicating bacteria promoting their survival includes ribosome hibernation (14, 15), which is characterized by programmed and reversible recruitment of specialized proteins to the decoding center of the ribosome (14). These proteins inactivate and preserve non-translating ribosomes as 70S monosomes or 100S disomes through mechanisms that differ across bacterial species (14, 16). In *Mycobacterium smegmatis* and Mtb, four distinct ribosome hibernation factors have been identified: Mpy, RafH, Balon and Rtof (17–21). While Mpy, RafH and Rtof hibernate a non-translating ribosome particle as a monosome, Balon is unique that it binds to the vacant A-site of a translating ribosome in the elongation complex (20). Rtof expression is induced *in vitro* in matured biofilms and leads to its binding to the nascent polypeptide exit tunnel (NPET). RafH expression is induced under hypoxia (18), although the proportion of ribosomes bound to RafH in hypoxic mycobacteria remains unknown. Mpy is constitutively expressed at high levels, but its association with the ribosome increases during limiting nutrients, including zinc (18, 19, 22). Zinc is an essential micronutrient for all organisms, and, like nutrient starvation or hypoxia, severe zinc limitation restricts mycobacterial growth. Zinc-limiting cultures of *M. smegmatis*, *M. tuberculosis and M. abscessus* have been observed to have high levels of Mpy-bound 70S ribosomes (16, 23). Mpy-dependent hibernation of 70S ribosomes is likely among the late stages of a multiphase zinc starvation response in mycobacteria (16) (Fig. 1A). Prior to hibernation, remodeling of the ribosome is induced, in which ribosomal proteins (r-proteins) containing the zinc binding CXXC motif (C+) are replaced by their C- paralogues lacking this motif (19, 24, 25). Ribosome remodeling is induced by transcriptional de-repression of the Zur-regulated C- r-gene operon, which occurs under zinc limiting conditions that are less severe than those inducing ribosome hibernation (16). The induced C- r-proteins replace the C+ protein paralogs and maintain translation activity of the ribosome, thereby supporting protein synthesis and growth in mycobacteria (16, 19, 24–27). A key factor facilitating the maintenance of stable Mpy-70S complex, called Mpy recruitment factor (Mrf), is transcriptionally co-induced with C- r-proteins (16, 28). Mrf is a zinc-binding protein, and its zinc-bound form is recognized and turned over by Clp protease during the expression of the C- ribosome. However, progressive depletion of intracellular zinc reduces Clp-mediated Mrf degradation, thereby stabilizing Mrf. Mrf then promotes the formation of a stable Mpy-70S complex (16).

**Figure 1:**
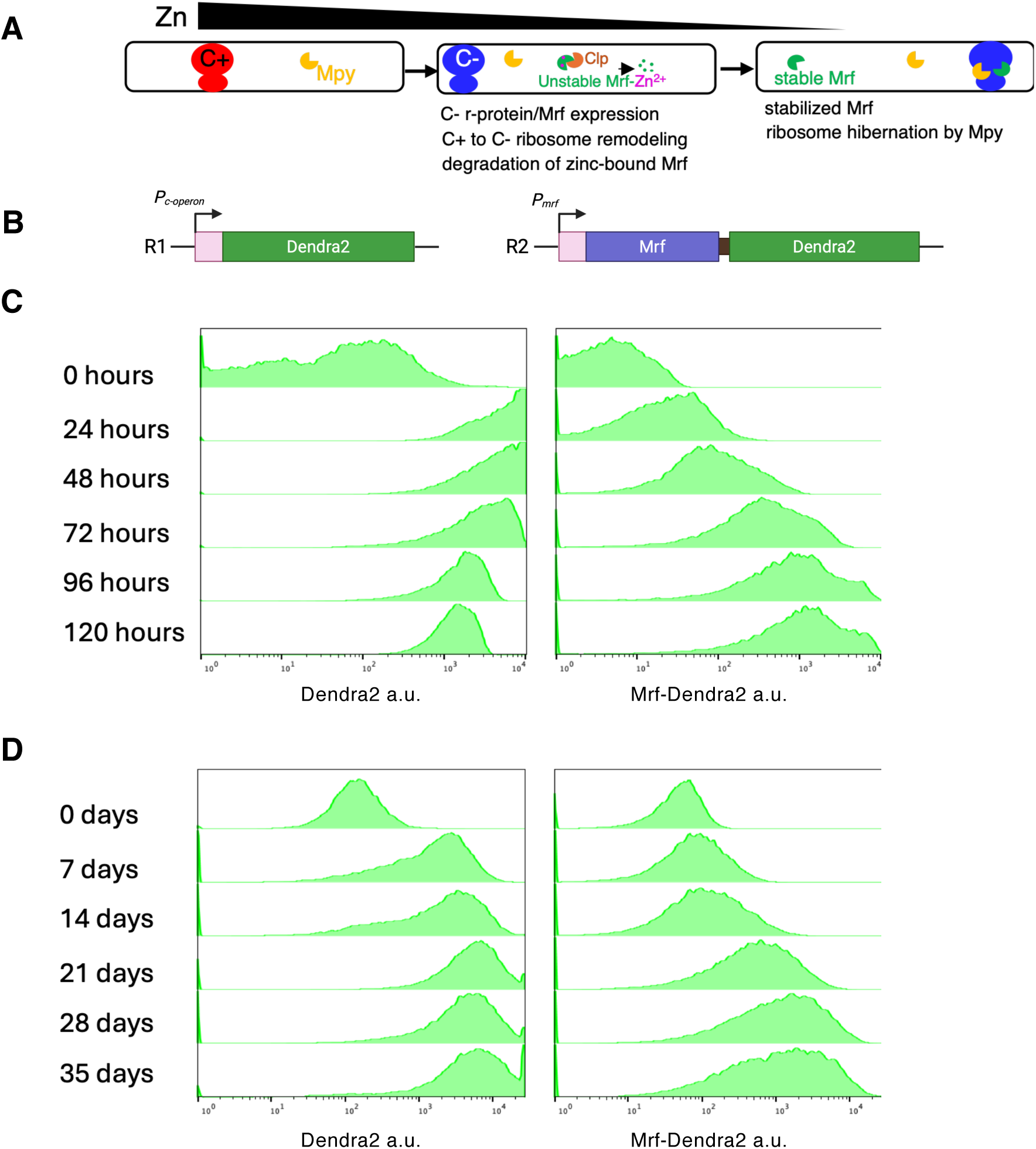
Reporters to visualize ribosome remodeling and Mrf accumulation as distinct zinc-starvation responses in mycobacteria. **A-B.** Zinc-responsive progressive changes in the ribosome from remodeling to hibernation through transcriptional de-repression of Zur-controlled C- r-proteins and Mrf, followed by post-translationally regulated stabilization of Mrf, which promotes Mpy recruitment to the remodeled (C-) ribosome. B. A schematic of fluorescent reporters for ribosome remodeling and Mrf stability, namely R1 and R2, respectively, are shown below. **C-D.** Flow-cytometric analysis of R1 and R2 reporters in *M. smegmatis* mc^2^155 (C), and Mtb mc^2^7000 strain (D). Cells were collected at the indicated time of growth (denoted on the left) in low-zinc Sauton’s medium and 20000 cells from each culture were analyzed by a flow cytometer. Additional characteristics of the R2 reporter are provided in supplementary figures S2-S5.

The clinical relevance of zinc-responsive Mpy-dependent ribosome hibernation is highlighted by: a) the evidence of zinc starvation response in Mtb in mouse lungs (19, 29) or in sputa of TB patients (30), b) Mpy-dependent reduced streptomycin sensitivity in Mtb in mouse lungs (16), and c) enrichment of Mtb expressing C- ribosomes in antibiotic-treated mouse lungs (31). Together, these studies suggest greater drug tolerance in zinc-starved subpopulations of Mtb *in vivo*. Although *in vivo* origins of zinc starvation in Mtb remain unknown, studies in other bacterial pathogens suggest that zinc starvation is induced by hosts as a component of the innate immune response called nutritional immunity (32). Zinc sequestration in extracellular niches of a pathogen is likely caused by a zinc-chelating heterodimeric complex composed of S100A8/A9 protein, called calprotectin (CP), and perhaps, to some extent, by a subset of plasma-membrane associated Zrt-Irt-like proteins (ZIP), which transport zinc into the cytoplasm (32, 33). Up to 40% of the cytoplasmic content of a neutrophil is CP (32), suggesting an important role of neutrophils in sequestering extracellular free zinc.

A high frequency of Mtb bacilli expressing C- r-genes was detected in chronic mouse lung infections (19). However, the proportion of these bacilli experiencing the extent of zinc starvation necessary to stabilize Mrf and form stable Mpy-70S complex has remained unknown. Exploiting the post-translational regulation of Mrf, we developed a translational fusion of Mrf with a fluorescent protein (Dendra2) reporter to visualize mycobacterial zinc starvation response that leads to stabilization of Mrf. The Mrf-Dendra2 reporter has allowed us to observe Mpy-dependent growth attenuation in Mtb at a single-cell resolution *in vitro*, overcoming the experimental challenges raised by cell-to-cell heterogeneity in the population. Mouse infection with Mrf-Dendra2 reporter strain showed that zinc-starved bacilli with high-levels of fluorescent signal, and presumably high levels of Mrf and Mpy-bound 70S ribosomes, constitute ∼3% of total bacterial population, and their frequency increases to ∼35% upon isoniazid treatment. Finally, investigation of host factors revealed that zinc mobilization by neutrophils plays an important role in the formation of this subset of bacilli.

## Results

### Quantification of remodeled and hibernating ribosomes in zinc-starved Mtb

To quantitatively assess the zinc-responsive changes in the ribosome, we purified 70S particles from cells of an attenuated strain of Mtb (mc^2^7000) cultured in high- and low-zinc Sauton’s medium, which previously differentiated the levels of Mpy in the ribosomes (16). Relative abundance of individual ribosomal proteins and Mpy in the ribosomes was determined by tandem-mass-tag mass spectrometry (TMT-MS). In low-zinc Mtb cells, relative abundance of S18c-, L28c- and L33c- was ∼6-fold higher than the median ratio of the invariant proteins, while their C+ counterparts were concomitantly reduced (Fig. S1). The L28_C-_ paralogue encoded by Rv0105c was not detected, confirming that Rv2058c encoded L28_C-_ is the primary constituent of the C- ribosome. The relative abundance of S14_C-_ was modestly increased (∼2-fold) in low-zinc ribosomes, but S14_C+_ abundance remained unchanged (Fig. S1). This suggests heterogeneity in the ribosomal composition of zinc-starved Mtb cells with respect to S14 proteins and thus, a possibility of chimeric 70S ribosomes assembled with S14_C+_ and S18_C-_, L28_C-_ and L33_C-_. The relative abundance of Mpy increased by ∼2-fold in C- ribosomes (Fig. S1), offering a measure of difference in the levels of Mpy-bound ribosome upon zinc depletion. While growth attenuation by limiting zinc is a reasonable condition for inducing ribosome hibernation, it does not rule out ribosome hibernation by Mpy or its paralogue Rv0079 by other growth-limiting conditions such as macronutrient starvation or hypoxia (18, 22).

### Visualizing mycobacteria with stabilized Mrf

We next sought to develop a method to visualize zinc-starved Mtb with a physiology that represents high-level of Mpy-bound ribosomes. A zinc-responsive accumulation of Mrf through a post-translational regulation induces Mpy recruitment to the ribosome (16, 28). We therefore rationalized that expression of a translational fusion of Mrf to a fluorescent protein Dendra2 from the native *mrf* promoter (*P_mrf_*-Mrf-Dendra2), henceforth called R2 reporter (Fig1A-B), would indirectly report cellular physiology that supports Mrf stability and Mpy-dependent ribosome hibernation. Due to the critical role of the N-terminal region of Mrf in its post-translational regulation (16), we fused Dendra2 to the Mrf C-terminus. Mrf and Dendra2 transcription was constitutive in a Δ*zur* strain harboring the R2 reporter (Fig. S2A), although Mrf-Dendra2 fluorescence signal was detectable only upon zinc depletion (Fig. S2B), indicating that fusion with Dendra2 doesn’t significantly impact the features of Mrf necessary for its zinc-dependent post-translational regulation. We note that Mrf-Dendra2 signal in the R2 reporter is merely a surrogate marker for a low-zinc physiology conducive for ribosome hibernation, because the reporter is functionally impaired. Unlike a functional Mrf-FLAG, multicopy constitutive expression of which inhibits growth (16), constitutive expression of Mrf-Dendra2 was well-tolerated by zinc-starved cells (Fig. S3).

The R2 reporter strains of *M. smegmatis* mc^2^155 and Mtb mc^2^7000 were cultured in low-zinc Sauton’s medium, and Mrf-Dendra2 signal was analyzed at various intervals by flow cytometry. A *P_mrf_*-Dendra2 reporter, called R1 (Fig. 1A-B), was used as a control to visualize the timing of Mrf transcription. Both reporter strains also harbored a plasmid expressing mCherry reporter (R3) from a constitutive *hsp60* promoter. Transcriptional de-repression of Dendra2 in the R1 reporter strains of *M. smegmatis* peaked after 24 hours, while the Mrf-Dendra2 fluorescence in the R2 reporter strain gradually increased to reach a saturating level by 96 hours (Fig. 1C), a timeframe by which high levels of Mpy-bound ribosomes were observed in the cell (Fig. S4A). This temporal separation between transcription of Mrf and accumulation of Mrf-Dendra2 was also observed in Mtb (Fig. 1D). Consistent with Dendra2 expression from R1 reporter (Fig. 1C), the reporter transcription in the R2 strain of *M. smegmatis* was induced by 24 hours (Fig. S4B), confirming that Mrf-Dendra2 accumulation is regulated at the post-translational level. In both species, the Mrf-Dendra2 signal in the R2 reporter population was distributed across a broad intensity range (Fig. 1C-D), indicating significant heterogeneity in the population. The heterogeneity could be visually confirmed microscopically for both *M. smegmatis* (Fig. S4C) and Mtb (S5A), based on which we defined Mrf-Dendra2-positive and negative cells as P1 and P2 subpopulations, respectively. We further noted that in contrast to *M. smegmatis* (Fig. S4C), the Mrf-Dendra2 signal in Mtb predominantly appeared as puncta (Fig. S5A). The puncta formation was not an artifact of fixing Mtb cells with paraformaldehyde (Fig. S5A), which has been shown to localize Dendra2 signal in mammalian cells (34). Moreover, Mrf-Dendra2 puncta in zinc-starved Mtb cells remained stable upon their subsequent exposures to the two host-relevant stressors, acidic pH and hypoxia (Fig. S5B-C).

### Relationship between Mrf-Dendra2 accumulation and growth of mycobacteria

To objectively evaluate Mrf-Dendra2 accumulation as an indirect visual marker for cells with hibernating ribosome, we tested the prediction that detectable fluorescent signal in P1 subpopulation of R2 reporter strain would correlate with two key phenotypes expected in cells harboring high levels of Mpy-bound ribosomes: slow metabolic activity and growth attenuation. Metabolic activities in P1 and P2 cells of R2 reporter of parental *M. smegmatis* mc^2^155 and isogenic Δ*mpy* or complemented strains were determined using 5-carboxytetramethylrhodamine amino-D-alanine (RADA), a fluorescent analogue of D-alanine. RADA is incorporated into newly synthesized peptidoglycan in growing cells (35). Relative to the P2 cells, RADA intensity was significantly lower in the P1 cells of mc^2^155, and this difference was abrogated in the Δ*mpy* strain, which also had overall higher RADA incorporation in both P1 and P2 cells (Fig. 2A). Complementation of Δ*mpy* with a plasmid-borne copy of *mpy* significantly reduced the RADA incorporation in both P1 and P2 cells (Fig. 2A), although the difference between P1 and P2 cells could not be recapitulated, perhaps due to partial complementation in which a larger difference in the mutant strain from the parent could be restored, but a relatively smaller difference between two subpopulations (P1 and P2) couldn’t be resolved. The weaker RADA staining in the parental P1 cells led us to predict that they would exhibit slow recovery in fresh nutrients relative to their P2 counterparts. When monitored by time-lapse microscopy in a microfluidic chamber perfused with zinc-limiting fresh M63 medium for 30 hours, P1 cells indeed exhibited retarded outgrowth compared to their P2 counterparts in a Mpy-dependent manner (Fig. 2B and supplementary movie 1A-B). The slower outgrowth of P1 than P2 cells was further corroborated by delayed uptake of RADA by P1 cells under this condition (Fig. S6A-B). However, this distinction between P1 and P2 cells appeared to originate from limiting nutrients and zinc in M63-TPEN medium, and possibly restricted oxygen supply in the microfluidic chamber, because both subpopulations outgrew to form colonies on nutrient-rich agar plates (Fig. S6C).

**Figure 2:**
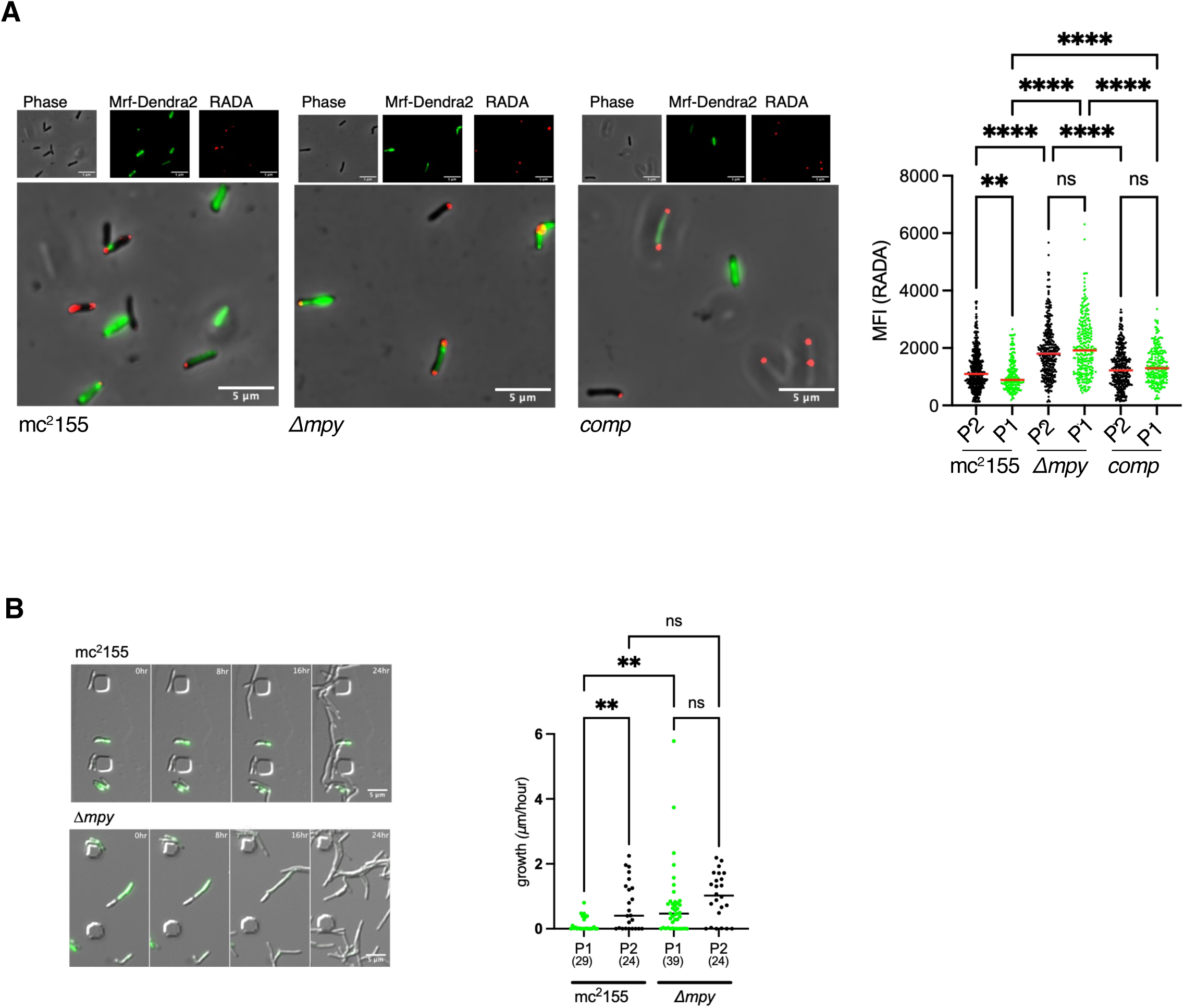
Mrf-Dendra2 marks Mpy-dependent metabolic quiescence in zinc-starved *M. smegmatis*. **A.** Representative micrographs showing differential RADA staining in P1 and P2 subpopulations of R2 reporter of parent (mc^2^155), *Δmpy* and *Δmpy* complemented (comp) strains of *M. smegmatis.* Plot shows RADA mean fluorescent intensity (MFI) at the poles of individual cells from three biologically independent experiments. **B.** Outgrowth of P1 and P2 cells of mc^2^155 and *Δmpy* in microfluidic chambers at indicated hours after initiating perfusion with M63 minimal medium containing 1 μM TPEN (also see supplementary figure S6A-C and movies S1A and S1B). Plot shows average elongation rate (μm/hr) of 24-39 (numbers for each sample indicated below) P1 and P2 cells of both strains from four biologically independent experiments. In panels A and B, ** and **** represent P < 0.01 and 0.0001, respectively (Unpaired Kruskal-Wallis ANOVA).‘ns’ denotes not significant.

The differential RADA staining offers crucial insights into the metabolic states of P1 and P2 cells; P1 cells appear to be in a slower metabolic state, presumably with high levels of Mpy-bound ribosomes, than P2 cells. Moreover, the greater RADA signal and accelerated growth in Δ*mpy* P1 cells suggest that Mpy-dependent ribosome hibernation and associated metabolic slowdown is induced prior to complete exhaustion of essential nutrients. Taken together, Mrf-Dendra2 accumulation in a cell expressing wild-type Mpy and Mrf can be reasonably inferred as an indirect marker for a slow-replicating state induced by ribosome hibernation, although the reporter *per se* cannot measure the level of Mpy-bound ribosomes in the cell.

### Impact of Mpy on physiology and isoniazid tolerance in Mtb

We next investigated the effect of Mpy-dependent ribosome hibernation on metabolic state of Mtb and its response to isoniazid (INH), a drug that selectively targets actively growing cells (36). Comparing RADA uptake in Mtb (mc^2^7000) R2 reporter cells with its isogenic Δ*mpy* mutant, with and without INH exposure, we observed that P1 cells with microscopically detectable levels of Mrf-Dendra2 incorporated lower levels of RADA than both its P2 counterparts, and P1 cells of Δ*mpy* strain (Fig. 3A). Treatment with INH significantly reduced the RADA incorporation in P2 cells of mc^2^7000, and P1 cells of Δ*mpy* mutant (Fig. 3A), suggesting that both these subpopulations are metabolically active enough to remain sensitive to INH. Consistent with the slow-metabolic state of mc^2^7000 P1 cells, their RADA signal did not change upon INH treatment (Fig. 3A). The Mpy-dependent INH sensitivity of Mtb was subsequently confirmed at the population level in a batch culture, in which starved Δ*mpy* cells had significantly fewer frequency of survivors from INH exposure than the parent mc^2^7000 and complemented strains (Fig. 3B). The MIC of the drug against the actively growing culture of the two strains was similar (Fig. 3C), suggesting that the minor subpopulation of survivors is likely exhibiting phenotypic resistance to the drug.

**Figure 3:**
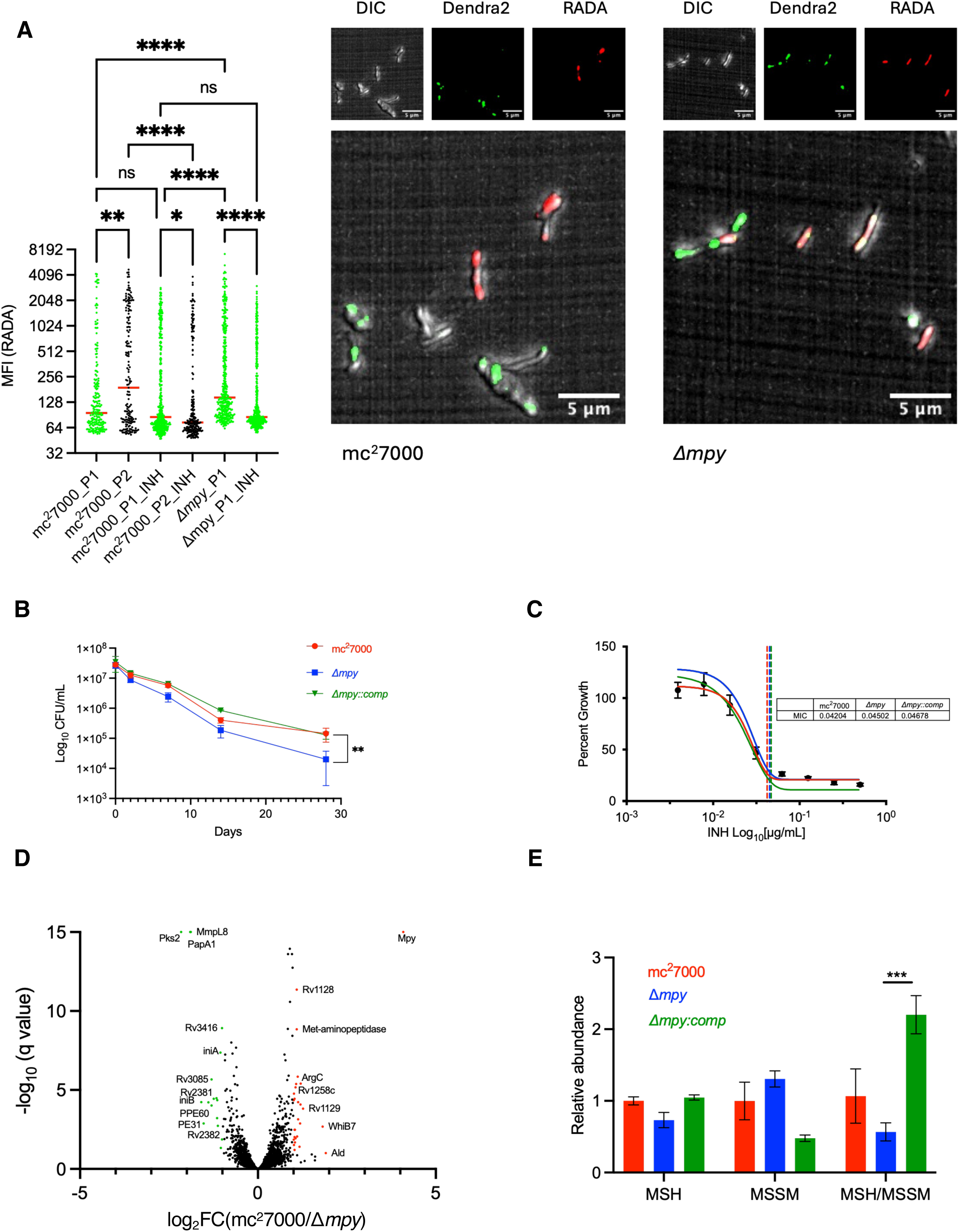
Mpy-dependent metabolic quiescence and INH tolerance in Mtb, marked by Mrf-Dendra2 accumulation. **A.** MFI of RADA at the poles of P1 and P2 cells of R2 reporter of parent Mtb strain (mc^2^7000) and *Δmpy* strains, either untreated or treated with ΙΝΗ. Cells were cultured in low-zinc Sauton’s media for 7 weeks, then exposed to 50 μg/mL INH for 7 days, after which an aliquot was washed and resuspended in 7H9OADCTw containing 50 μM RADA, incubated overnight at 37 °C prior to visualization under microscope. Also shown are the representative micrographs of each strain. **B.** Survival of Mtb mc^2^7000, *Δmpy*, and complemented (comp) strains after exposure to INH. Cells were cultured for 7-weeks in low-zinc Sauton’s medium, then suspended in PBST with 50 μg/mL INH and plated at indicated timepoints after exposure. **C.** MIC of INH for Mtb mc^2^7000, *Δmpy*, and complemented (comp) in low-zinc Sauton’s medium. The lines represent MIC_90_ the values of which are indicated in the table. **D.** RNA-seq-derived volcano plot showing differentially expressed genes between mc^2^7000 and *Δmpy* strains cultured in low-zinc Sauton’s medium (full dataset is available at GSE263900). Significantly (q-value < 0.05) up- and downregulated genes by > 2-fold are marked with red and green colors, respectively. **E.** Relative levels of mycothiol to mycothione ratio determined through metabolome profiling by LC-MS/MS of Mtb mc^2^7000, *Δmpy* and complemented (comp) strains cultured for 7 weeks in low-zinc Sauton’s medium. Values indicate average of three biologically independent experiments normalized to glutamic acid as an unaffected internal control (also see supplementary figure S7). *, **, *** and **** represent P < 0.05, 0.01, 0.001 and 0.0001, respectively (Unpaired Kruskal-Wallis ANOVA for panel A, t-test for panel C and F).

To further understand the impact of Mpy on Mtb progression to dormancy, we compared the transcriptome of mc^2^7000 and Δ*mpy* strains, with a focus on redox-sensitive genes. We reasoned that the slowed metabolism from Mpy-dependent ribosome hibernation would lower the production of endogenous reactive oxygen species, thereby impacting the intracellular redox environment, which in turn would alter the expression of redox-sensitive genes. An RNA-seq based transcriptional analysis of low-zinc cultures of mc^2^7000 and Δ*mpy* strains identified 41 genes with significantly (>2-fold; q < 0.05) different expression between the two strains (Fig. 3D; complete dataset at Gene Expression Omnibus GSE263900). Among the 16 genes with elevated expression in Δ*mpy* cells were *pks2*, *papA1* and *mmpL8*, organized in an operon. Pks2 is upregulated by WhiB3, a redox sensing transcriptional regulator that binds and activates the *pks2* promoter upon oxidation of its cysteine thiol groups (37). Prominent among the 25 downregulated genes in Δ*mpy* was *whiB7* (38) (Fig. 3D, GSE263900). The Mpy-responsive expression of WhiB7 was further consistent with downregulation of multiple amino acid biosynthetic genes including AspC and ArgBCD (see GSE263900), which have been previously shown to be activated by WhiB7 (39, 40). Unlike WhiB3, WhiB7 expression is induced by a thiol-reducing agent like DTT and correlates with high mycothiol/mycothione (MSH/MSSM) ratio (41). Thus, the opposing expression pattern of WhiB7 and WhiB3-dependent genes reflect an elevated level of endogenous intracellular oxidative stress in a Δ*mpy* strain than in the parent mc^2^7000 strain. A lower MSH/MSSM ratio in the Δ*mpy* strain, compared to the parent and complemented strains, was confirmed by profiling the whole-cell metabolites (Fig. 3E, Fig. S7).

Mpy-bound ribosomes are directly protected from streptomycin and kanamycin (19). However, the evidence of Mpy-dependent WhiB7 expression, which induces resistance to multiple ribosome-targeting drugs (38), and redox homeostasis that determines INH sensitivity (36), suggest that Mpy may have a broader impact on drug tolerance in Mtb through indirect mechanisms. Mpy-dependent INH tolerance could likely emerge from ribosome hibernation and metabolic slowdown, resulting in relatively low levels of intracellular reactive oxygen species (ROS) usually generated as byproducts of aerobic respiration (36). We note that genes induced by INH exposure in Mtb, *iniA*-C (42), are also induced in unexposed Δ*mpy* strain (Fig. 3D), suggesting a shared physiological stress between the mutant and INH-exposed cells.

### Visualizing isoniazid-tolerant P1 cells of Mtb in mouse lungs

Our *in vitro* study thus far identifies Mrf-Dendra2 as a suitable reporter for visualizing slow-replicating mycobacteria under severe zinc starvation. We next asked if zinc-concentrations in host tissues are low enough to give rise to P1 subpopulation of Mtb *in vivo*, and if so, then how this subpopulation is impacted upon treatment of the infection with INH. Lung sections from C3HeB/FeJ mice, infected for 8-weeks with Mtb(Erd):R2:R3 reporter strain and subsequently treated with INH for 2- or 4-weeks, were microscopically visualized. The C3HeB/FeJ strain was chosen because Mtb infection in this strain produces a neutrophil-rich human-like lung pathology with high bacterial burden that are difficult to treat with antibiotics (4, 43). The 8-week timepoint represents the chronic phase of infection when a majority of Mtb bacilli in C3HeB/FeJ mice express C- ribosomes (19). Approximately 3% of the total bacterial population in the pre-treated group accumulated detectable levels of Mrf-Dendra2, and the frequency increased progressively during the course of INH treatment in an Mpy-dependent manner (Fig. 4A-D, Fig. S8). The increase in the frequency is most likely due to preferential killing of the P2 subpopulation without the signal. Interestingly, the difference in the frequency of INH survivors among P1 subpopulations of WT and *mpy* mutant does not impact the overall bacterial burden in the lung within a 4-week treatment period (Fig. 4C-D). An explanation can be obtained from analyzing the rate of clearance of P1 and P2 subpopulation, which decline by ∼75% and 99.2%, respectively, over a 4-week INH treatment. Thus, while majority of P1 cells *in vivo* remain susceptible to INH, their survivor frequency remains higher than in P2. Moreover, in a hypothetical situation, even if P1 cells were to reduce similarly as P2 cells, the gross bacterial burden after 4-weeks of INH treatment would reduce only by about 32%, which is too small a difference to be distinguished by the serial dilution method.

**Figure 4:**
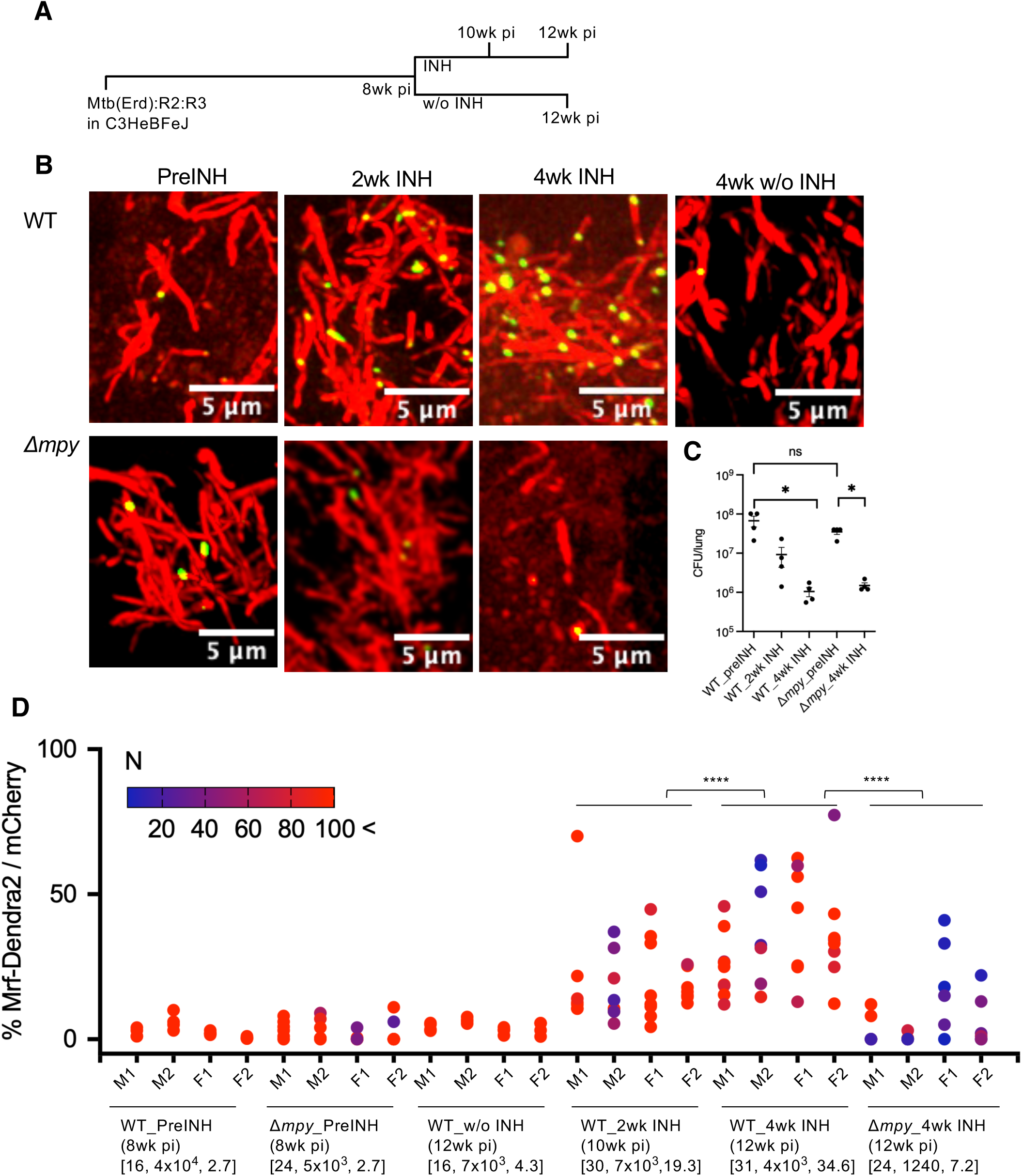
Visualization of Mtb(Erd) accumulating Mrf-Dendra2 in mouse lungs. **A.** An overview of the experimental design in C3HeBFeJ mouse strain. Mice were infected for 8-weeks with R2 reporter strain of wild-type Mtb(Erd) (denoted WT), or its isogenic *Δmpy* mutant. Both strains constitutively expressed mCherry (reporter R3). A group of 4 mice each was treated with INH for either 2 or 4 weeks, while another group was left untreated for 4 weeks. Lung sections at indicated timepoints were microscopically visualized. **B.** Representative micrographs of lung sections show merged maximum projections of red- and green channels. Colocalization of Mrf-Dendra2 signal with mCherry was confirmed as shown in supplementary figure S8. **C.** Gross bacterial burden in the lungs of above-described infection before and after INH treatment. **D.** A multi-variable plot showing frequency of Mtb bacilli accumulating Mrf-Dendra2 before and after INH treatment of male (M1, M2) and female (F1, F2) mice. Each dot represents a field-of view (FOV), and its color represents the number of bacilli visualized (saturation limit of 100). The three values below each infection group denote aggregate data from four mice: (from left) total number of FOV analyzed, total number of bacilli visualized, and the mean frequency of bacilli with Mrf-Dendra2 signal. *, **** denote P < 0.05 and 0.0001, respectively (Mann-Whitney).

To gain insights into the host environment associated with Mtb P1 subpopulation, we visualized bacilli and neutrophils in lung sections from mice infected for 8-weeks with Mtb(Erd):R2:R3 reporter strain. Our approach was based on the premise that P1 bacilli would be detected in lesions with degranulating neutrophils releasing CP (32, 33, 44). In a preliminary analysis, P1 subpopulation was often found in distinct clusters of bacilli that were surrounded by Ly6G-expressing neutrophils (Fig. 5A). Moreover, lack of DAPI signal around such bacterial clusters suggested that these perhaps developed within a necrotic lesion (Fig. 5A).

**Figure 5.**
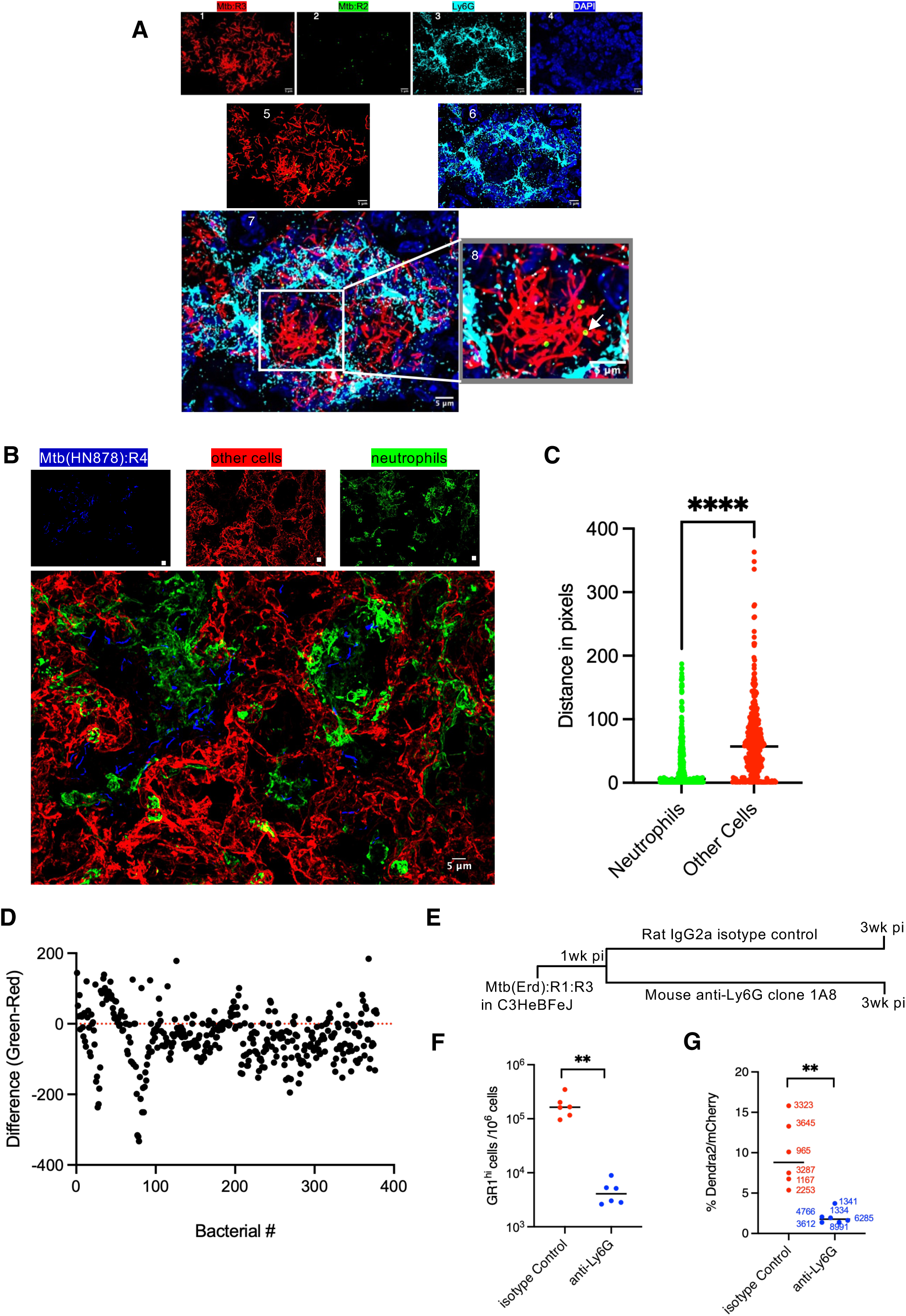
Neutrophils induce zinc starvation response in Mtb. **A.** Immunofluorescence staining with anti-Ly6G antibody of a C3HeBFeJ mouse lung section infected for 8 weeks with WT Mtb(Erd):R2:R3. Individual channels of red (1; all Mtb bacilli expressing mcherry), green (2; Mtb accumulating Mrf-Dendra2), cyan (3; anti-Ly6G) and blue (4; DAPI) are shown at the top, while the merged channels of Mtb signals (5; red + green) and host signals (6; cyan + blue) are shown in the middle. All channels are merged at the bottom in 7. The rectangle (magnified in 8) indicates a likely necrotic center harboring bacilli surrounded by neutrophils, including few with Mrf-Dendra2 signal. **B.** Visualization of Mtb(HN878) expressing BFP from C- r-protein promoter (blue), Mtb(HN878):R4, in the spatial context of neutrophils (expressing EGFP) and other cells (expressing tdTomato) in a reporter mouse S100a8-Cre ^mT/mG^ (also see supplementary figure S9). Thumbnails on the top show individual color channels. **C-D.** Distance measurement from an individual bacterial cells (blue) to the nearest neutrophil (green) or nearest non-neutrophil (red) cell. Plot in panel C shows overall proximity distribution from a bacillus to the nearest neutrophil (green) and non-neutrophil (red) cells [n= 376, **** denotes P(t-test) < 0.0001]. Plot in panel D shows the difference in the distance from a given bacillus to a green and red cells. Approximately 73% of 376 analyzed bacilli were closer to neutrophils than to non-neutrophil cells. Data were collected from five sections across two infected mice. **E-G.** A causal relationship between neutrophils and zinc starvation response in Mtb. Panel E shows an overview of the experimental design. Mtb(Erd):R1:R3 reporter strain expressed Dendra2 from the C- r-protein promoter (R1) and mCherry from the constitutive *hsp60* promoter (R3). Frequencies of neutrophils (GR1^hi^ cells) and Dendra2 expressing Mtb cells in each mouse lung are summarized in panels F and G, respectively. Each datapoint in both panels represent one mouse. Datapoint in panel G is obtained from indicated number of bacilli from multiple lung section of a mouse. Additional flow cytometry data, including the gating strategy, micrographs and lung bacterial burden are provided in supplementary figure S10. ** denotes P < 0.01 (Mann-Whitney).

In an alternative and more comprehensive approach to visualizing the spatial relationship between neutrophils and zinc-starved Mtb, we generated a transgenic mouse strain in C57BL/6 background that expressed GFP in the membrane of neutrophils and tdTomato in the membrane of other cells (Fig. S9A-D). This mouse strain was infected with a reporter strain of Mtb(HN878), called Mtb(HN878):R4, which expressed blue fluorescent protein (BFP) from the C- r-protein promoter under zinc limiting condition (Fig. S9E). HN878, a hypervirulent Mtb strain, causes extensive necrotic lesions even in a relatively more resistant C57BL/6 strain (45), thereby allowing easy visualization of neutrophils. Expectedly, ∼70% of BFP-expressing bacilli were closer to neutrophils than non-neutrophil cells (Fig. 5B-D), indicating that zinc-starved Mtb bacilli are predominantly located in neutrophil-rich lesions.

For a direct causal relationship between neutrophils and zinc starvation response in Mtb, we tested the effect of antibody-mediated neutrophil depletion on the frequency of Mtb cells inducing C- ribosome expression. C3HeB/FeJ mice were infected with Mtb(Erd):R1:R3 reporter strain expressing Dendra2 from the C- r-protein promoter, and neutrophils were depleted by intraperitoneal administration of anti-Ly6G antibody (46). Immunophenotyping confirmed neutrophil depletion upon anti-Ly6G treatment (Fig. S10), which led to a significantly reduced frequency of Dendra2-expressing Mtb cells (Fig. 5E-G), indicating that zinc starvation response in Mtb is primarily induced by neutrophils.

### Expression of zinc mobilizing factors in neutrophils upon Mtb infection

Neutrophil infiltration is strongly associated with progressive pulmonary TB pathology (47–50), predicting that differential lung neutrophil abundance between the resistant and susceptible hosts like C57BL/6 and C3HeB/FeJ mice (51), respectively, could impact the proportion of P1 bacilli. Relative to C3HeB/FeJ mice, chronic infection of C57BL/6 mice with Mtb:R2:R3 strain expectedly produced less inflammation, fewer lung neutrophils, lower lung bacterial burden, and lower frequency of P1 bacilli (Fig. 6A-D). To further identify the host factors that could influence the development of P1 bacilli, we analyzed the differences in the host transcriptional response to chronic infection of Mtb(Erd):R2:R3 strain in C57BL/6 and C3HeB/FeJ mice, using uninfected mice as controls. Because of the relationship between neutrophils and zinc starvation response in Mtb, we prioritized our analysis to genes involved in zinc mobilization (ZIP, ZnT and S100A8/A9) and immune/inflammatory responses (GO0009654-55). Genes associated with the recruitment of monocytes (Ccl2, Ccl3, Ccl4, Ccl7, Ccl8, Ccl12) and neutrophils (Cxcl1, Cxcl2, Cxcl3, Cxcl5) were upregulated in C3HeB/FeJ mice compared to C57BL/6 (Fig. 6E, see GSE263940 for complete dataset). Moreover, the induced expression of a broader spectrum of inflammation-related genes (Nos2, Nlrp3, Ptgs2, Mefv, Tnf) were observed in the lungs of C3HeB/FeJ mice (Fig. 6E and GSE263940), consistent with the histopathological indication of tissue inflammation in figure 6A.

**Figure 6.**
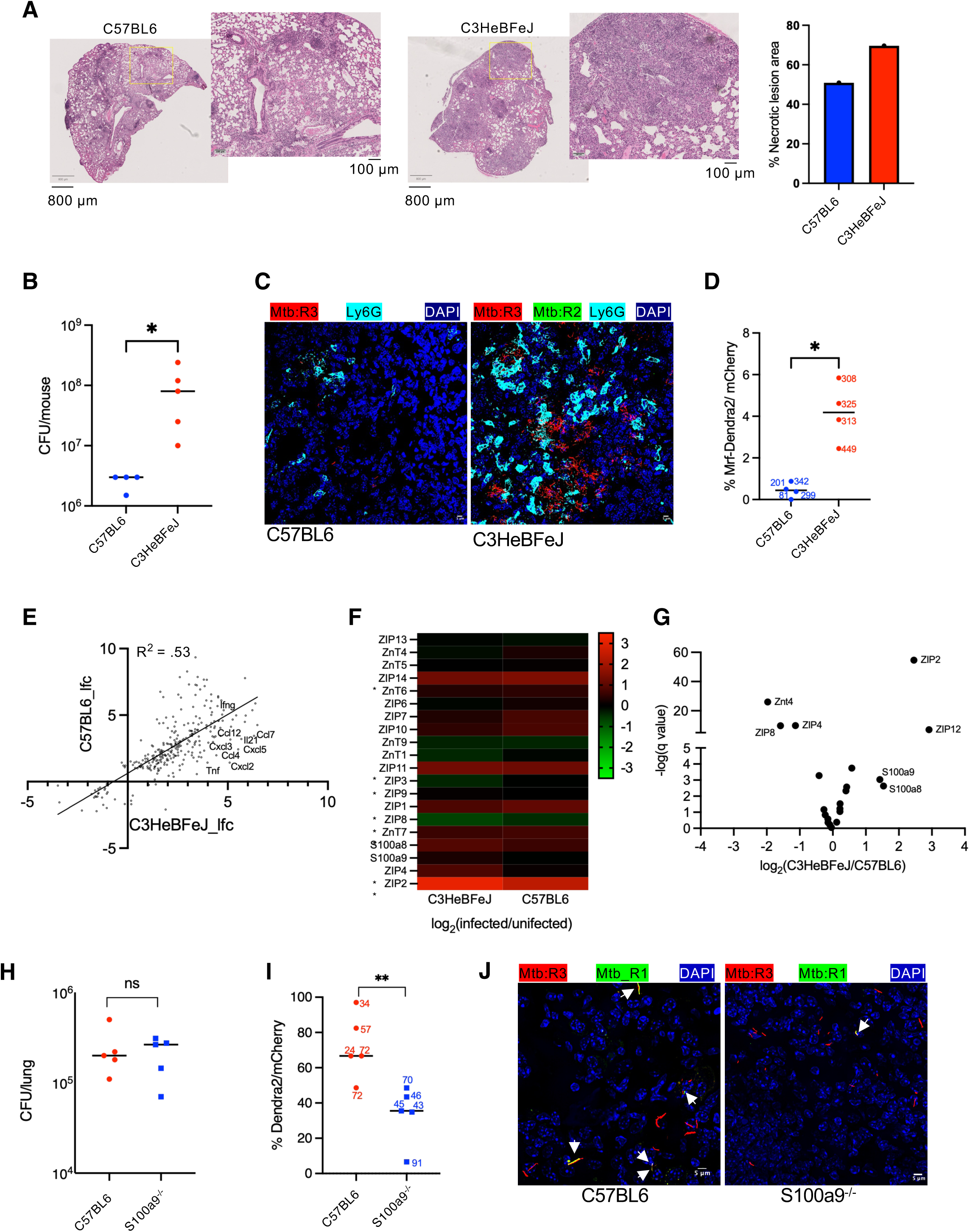
Calprotectin induces zinc starvation response in Mtb. **A.** Slide scan of H&E-stained lung sections of C3HeBFeJ and C57BL6 mice infected with Mtb(Erd):R2:R3 reporter strain for 12- and 14-weeks, respectively. These different timepoints were chosen to reflect late-stage chronic phase of infection, which is distinct for the two mouse strains. The adjacent micrograph corresponding to the inset is magnified to show greater tissue pathology in C3HeBFeJ than C57BL6 mouse. **B-D.** Correlation between bacterial burden (B), neutrophil abundance in lungs visualized with AF647-conjugated anti-Ly6G antibody (C), and the frequency of Mrf-Dendra2 accumulating Mtb (D), compared between C57BL6 and C3HeBFeJ mouse strains infected with Mtb(Erd):R2:R3 for 14- or 12-weeks, respectively. Each datapoint in panels B and D is an average value from one mouse. At least 4 sections per mouse were analyzed for panel D data and the numbers indicate the total numbers of bacilli analyzed. **E.** Relative transcript abundance of 280 inflammatory and immune response genes (GO: 0006954-55) that are significantly altered in lungs of either C3HeBFeJ or C57BL6 mice, or both, upon Mtb(Erd):R2:R3 reporter infection for 12- or 14 weeks, respectively. Significant (q-value < 0.05) log-fold-change (lfc) in the transcripts of each mouse strain was determined by lung RNA-seq from two uninfected and three infected mice, followed by DESeq2 analysis. **F.** Relative change in the expression of zinc transporters (ZIP and Znt encoding proteins) and two calprotectin chains (S100A8 and A9) from the same RNA-seq dataset described in panel E. * denote statistically significant (q-value < 0.05) change in expression upon infection. **G.** Volcano plot showing the relative difference in the expression of the panel F genes between neutrophils of C3HeBFeJ and C57BL6 mice, infected for 28 days with Mtb(HN878). Neutrophils from three mice of each strain were processed for RNA-seq. **H-J.** S100A9 deletion in C57BL6 mice reduces zinc-starvation response in Mtb. C57BL6 and isogenic S100A9^-/-^ deletion strains were infected with the reporter strain Mtb(Erd):R1:R3 for 14-weeks and total bacterial burden (H) and the frequency of Dendra2 expressing bacilli (I) were determined. Each datapoint in panel I is obtained from cumulative number of bacilli (as indicated) visualized in multiple lung sections of each mouse. Panel J shows representative micrographs from each mouse strain, in which bacilli expressing Dendra2 are indicated by arrows. * and ** in panels D and I denote P < 0.05 and 0.01, respectively (Mann-Whitney). ns denotes not significant.

With the exception of ZIP2, the expression of ZIPs and ZnTs as well as S100A8/A9 was modestly altered upon infection in either of the two mouse strains (Fig. 6F and GSE263940). ZIP2 expression interestingly was differentially induced in the two mouse strains: ∼8.5- and 5.2-fold in C3HeB/FeJ and C57BL/6 mice, respectively. ZIP2 predominantly transports zinc into the cytosol across the plasma membrane (52), suggesting that an induced ZIP2-dependent zinc influx into the cytoplasm concomitantly reduces the extracellular concentration, the predominant niche for the zinc-starved Mtb. ZIP8, which is induced in macrophages upon Mtb infection (53), was downregulated in the infected lung tissue of both mouse strains, although the effect was more pronounced in the C3HeB/FeJ lungs. The change in the expression of S100A8/A9 between C57BL/6 and C3HeB/FeJ mice was unexpectedly indistinguishable (Fig. 6F). Perhaps an upregulation of S100A8/A9 in neutrophils is neutralized by its negative regulation in other cells. We therefore analyzed the transcriptomes of lung neutrophils isolated from chronic Mtb infections in C57BL/6 and C3HeB/FeJ strains. For maximizing neutrophil influx in C57BL/6, we used the hypervirulent Mtb (HN878) (45). The S100A8/A9 expression in C3HeB/FeJ neutrophils was ∼3-fold higher than those from C57BL/6 counterparts (Fig. 6G and GSE244230). Somewhat consistent with the whole-lung-RNAseq profile, the ZIP2 expression between the two mouse strains was more pronounced than S100A8/A9 expression (Fig. 6G). In sum, S100A8/A9 and ZIP2 are likely the key contributors to zinc-limiting extracellular environment. To further determine the contribution of S100A8/A9 in neutrophil-induced zinc starvation response in Mtb, we compared the frequency of Dendra2 expressing cells of Mtb(Erd):R1:R3 reporter strain in C57BL/6 and isogenic *S100a9^-/-^* during the chronic phase (14-week) of infection. Partial but significant decrease in the frequency of Dendra2-expressing bacilli was observed in *S100a9^-/-^* strain (Fig. 6H-J), suggesting that the expression of additional factors like ZIP2 is necessary for optimum zinc starvation response in Mtb.

## Discussion

Phenotypic drug tolerance in mycobacterial population, first hypothesized in Mtb cells by Hobby and Lenerts eight decades ago (54), is a well-studied phenomenon under *in vitro* conditions (55–58). However, its occurrence *in vivo* remains untested. Here we describe an approach to visualize phenotypic isoniazid tolerance in Mtb bacilli (P1 cells) that develops through accumulation of Mrf, and likely Mpy-bound ribosomes, under stresses that include zinc-limiting host lung environment predominantly produced by infiltrating neutrophils. Neutrophils generate a variety of stresses for an invading pathogen, including oxidative damage and tissue hypoxia (59), which in addition to zinc starvation could possibly restrict growth and induce persistence through other mechanisms. Moreover, given that fluorescent signals in P1 cells persist under hypoxia and low pH, it is possible that INH tolerant subpopulation of P1 cells originate in response to a combination of stresses, while zinc-limiting environment is simply facilitating their visualization. Moreover, other types of INH tolerant cells not captured by this reporter likely exist in the lungs. However, prevalence of severely zinc-starved Mtb *in vivo* is unambiguous. Such level of zinc starvation probably emerges from the expression of ZIP2 and S100A8/9 in neutrophils, thereby sequestering zinc from extracellular Mtb by both chelation (S100A8/A9) and mobilization (ZIP2). Based on our *in vitro* evidence, we further infer that zinc-starved P1 cells of Mtb visualized by Mrf-Dendra2 acquire a slow-replicating state, likely due to high levels of Mpy-bound hibernating ribosomes. Their low frequency (∼3%), relative to the frequency of bacilli inducing ribosome remodeling (∼80-100%) (19), suggests a non-uniform distribution of free zinc in lungs that infrequently drops below the threshold necessary to induce Mrf stabilization in Mtb. This variation in free zinc levels, possibly due to the combined effects of neutrophils and interbacterial competition for zinc, could potentially be an important source of phenotypic heterogeneity among bacilli, although variations in the expression of other genes could further compound the effect.

The difference in the frequency of R1 (remodeling) and R2 (hibernation) reporters of Mtb, both *in vitro* and in mouse lungs, is consistent with the previously observed stepwise de-repression of the Zur regulon observed in *Bacillus subtilis* (60). C- r-protein expression is derepressed before that of zinc transporters, ZnuABC, likely due to progressive depletion of zinc that gives rise to partially metalated Zur dimer intermediates with varying levels of affinity for the promoters of *c-* genes and *znuABC*. Thus, different expression levels of zinc transporters, even among cells with remodeled ribosomes, can create a population-wide heterogeneity with respect to intracellular free zinc.

Downregulation of growth and metabolism as a broader outcome of Mpy-dependent translational control is a likely cause for antibiotic tolerance in Mtb cells, although the drug sensitivity of Δ*mpy* strains could also be compounded by other collateral effects of hypertranslation. One possible example is the frequent pausing of elongation complexes in the mutant. Mpy binding to the ribosome under nutrient-limiting conditions can arguably reduce the frequency of translation initiation to balance the pool of translating ribosome with other canonical protein synthesis ligands and amino acids.

The unexpected increase of WhiB7 expression in Mtb cells inducing ribosome hibernation is likely an indirect consequence of an overall reduction in translation by Mpy. WhiB7 transcription is typically induced by exposure to certain ribosome-targeting drugs through a mechanism that involves translation attenuation of an upstream open reading frame (uORF), such that ribosome inhibition or amino acid starvation allows transcription readthrough of the *whiB7* ORF (39, 61, 62). A slower rate of translation initiation on uORF in cells with high levels of hibernating ribosomes likely mimics the condition of antibiotic-mediated ribosome inhibition that leads to increased WhiB7 transcription. In addition, the increased WhiB7 transcription could also be an indirect outcome of the Mpy-dependent change in the intracellular redox homeostasis, which directly impacts the transcriptional auto-activation of WhiB7 (41). Given the emerging role of WhiB7 in positive regulation of amino acid biosynthesis (39), its upregulation in an Mpy-dependent manner could potentially boost the amino acid pool in bacilli. Thus, a possible scenario of ribosomal pausing during translation elongation in nutrient-starved bacilli can be minimized by two potentially additive Mpy-dependent mechanisms: by reducing the level of active ribosomes, and by inducing amino acid biosynthesis through induced WhiB7 expression. Through these mechanisms, the process of Mpy-dependent ribosome hibernation likely confers short-term adaptive advantages to bacilli growing under limiting-nutrients, while the stability of Mpy-bound intact non-translating ribosome in slow-replicating cells could serve as a long-term adaptation mechanism (19). In summary, these physiological insights into zinc-starved, slow-replicating population of Mtb, likely with hibernating ribosomes, along with the ability to visualize them in host tissues offer an experimental framework to understand and target these bacilli towards a shorter drug regimen for tuberculosis.

## Materials and Methods

### Bacterial growth medium

*M. smegmatis* (mc^2^155) strains were routinely cultured in Middlebrook 7H9 + 0.5% glycerol + 10%ADC + 0.05% Tween-80. Unless specified, *M. tuberculosis (*mc^2^7000) strains were routinely cultured in Middlebrook and 7H9 + 10%OADC + 0.5% glycerol + 0.05% Tween-80, Pantothenate (100 μg/mL). For culturing in high- and low-zinc Sauton’s medium, *M. smegmatis* cells from a 7H9ADCTw culture were centrifuged for 10 minutes using a Sorvall Legend XTR centrifuge with Thermo Scientific rotor 75003180 at 4000 rpm, washed 3 times with PBS + 0.05% Tween-80 (PBST), and inoculated in 50mL of Sauton’s medium + 0.05% Tween-80 supplemented with either 1mM ZnSO_4_ for high-zinc culture, or 1μM of TPEN [N,N,N,N’-Tetrakis (2-pyridylmethyl)ethylenediamine] as the zinc selective chelator for low-zinc culture.

Pellicle biofilms of *M. tuberculosis* (mc^2^7000) were cultured as described previously in detergent-free Sauton’s medium containing pantothenate (100 μg/mL) and supplemented with 1mM ZnSO_4_ for high-zinc condition or used without supplemental zinc for low-zinc condition. Cells from starter 7H9 +OADC+ 0.5% glycerol+ 0.05% Tween-80 +100 μg/mL PAN log phase planktonic cultures were inoculated at a 1:100 dilution into 4 mL of detergent-free Sauton’s medium in 12-well polystyrene plates and incubated without shaking for 7 weeks at 37 °C in humidified conditions.

### Reporter construction

Reporter 1 (R1) in *M. smegmatis* was constructed by cloning the promoter for Msmeg_6068 into the Xbal and BamHI sites of pYL026(16), an episomal plasmid expressing Dendra2 with hygromycin resistance marker (sequence in Table S2). R1 reporter for *M. tuberculosis* was constructed by cloning the promoter for Rv_2058c into the Xbal and NdeI sites of pYL027. Reporter 2 (R2) in *M. smegmatis* was constructed by cloning the native Msmeg_6069 with its native promoter into the XbaI and NdeI of pJL37 to create pYL164, and then a glycine rich linker sequence followed by Dendra2 at BamHI and HindIII into pYL164 (sequence in Table S2). R2 in *M. tuberculosis* was constructed by cloning a glycine rich linker sequence followed by Dendra2 into HindIII and NheI into pJL37 to create pYL170, and then Rv_0106 with its native promoter into XbaI and HindII into pYL170 (sequence in Table S2). Blue fluorescent protein (BFP) reporter was cloned into pYL001 by replacing Dendra2 at BamHI and HindIII sites.

### Microscopic analysis of reporter cells from in vitro batch cultures

*M. smegmatis*, reporter strains R1 and R2 were cultured in low- and high-zinc Sauton’s medium. For the first 36 hours, the OD600 was measured every 3 hours and cells were visualized using epifluorescent microscopy. After 36 hours, cells were collected once every 8 hours until 72 hours, and then once every 24 hours until 127 hours post inoculation.

For *M. tuberculosis*, cells were grown as biofilms for a total of 9 weeks in low- and high-zinc Sauton’s medium in 12-well plates. Starting at week 5, cells from one well were collected in a 15mL conical tube, centrifuged for 10 minutes in a Sorvall Legend XTR with Thermo Scientific rotor (#75003180) at 4000rpm, resuspended in PBS + 0.05% Tween-80, and dispersed overnight using 3 mm glass beads (Sigma, #Z143928-1EA) at 4 °C on a platform shaker. After dispersion, cells were centrifuged at 4000rpm for 10 minutes and fixed for 1 hour in 4% paraformaldehyde.

To prepare samples for microscopic analysis, 0.14mm microscope coverslips (Globe Scientific, #119-1401-10) were incubated with 200 μL of 1 mg/mL Poly-D-lysine hydrobromide, molecular weight 30,000-70,000 (Sigma, #45-P7886-50MG) for 5 minutes. The solution was then aspirated, and the coverslips were washed with sterile water and dried at room temperature. To mount the slides, 10 μL of Molecular Probes™ SlowFade™ Diamond Antifade Mountant (Thermo Fisher, #S36967) was applied to the slide, while 2.5μL of cell suspension was gently spread along the surface of the coverslip. The coverslip was then placed onto the slide with mounting media; excess media was removed using a Kimwipe. Slides were visualized immediately using a Nikon Eclipse TE2000-E with the X-Cite120 Fluorescence Illumination System with 60X and 1.4 N.A. objective. Images were captured using a Roper HQ camera and the ImagePro software using 700 millisecond preview exposure and 700 millisecond acquisition exposure with 2X2 binning for both the preview and acquisition exposures. Between 3 to 5 fields were acquired per sample. Individual cells were counted and analyzed using Fiji software. All cells R1 constitutively expressed mCherry from pEM2 and R2 cells from pJC009 in *M. smegmatis*. In *M. tuberculosis,* all R1 cells constitutively express mCherry from pEM2 and R2 cells from pYL192. The percentage of Dendra2 positive cells was calculated from the ratio of Dendra2 positive cells to mCherry positive cells.

### Phenotyping of reporter cells by flow cytometry

*M. smegmatis*, reporter strains R1 or R2 were cultured in low- or high zinc Sauton’s medium. Every 24 hours, 500 μL from each culture was diluted in 4 mL of PBS and fluorescent phenotyping was performed using a Becton Dickinson FACSCalibur flow cytometer. Cells from a total of 50,000 events were gated to exclude cellular debris and aggregates. Acquisitions were analyzed using FlowJo™ v10.8.2 software (BD Life Sciences). *M. tuberculosis* mc^2^7000 reporter strains, R1 and R2, were cultured for 1 week in Middlebrook 7H9 + 10%OADC + 0.5% glycerol + 0.05% Tween-80 + 100 μg/mL PAN until mid-log phase. Cells were then washed and cultured in Sauton’s + 100 μg/mL PAN without supplemental zinc (low-zinc) or with 1 mM ZnSO_4_ (high-zinc). After two weeks of initial growth in low-zinc Sauton’s media, 1 μM TPEN was added, and cells were grown for an additional three weeks. Starting when the cells were first cultured in Sauton’s media, a 1 mL aliquot of cells was retained and fixed in 4% paraformaldehyde in PBS once per week to monitor changes in fluorescent reporter expression. Fixed cells were diluted in 4 mL of PBS, and fluorescent phenotyping was performed using a Becton Dickinson FACSymphony™ A3 flow cytometer. A total of 20,000 events were gated to exclude cellular debris and aggregates. Acquisitions were analyzed using FlowJo™ v10.8.2 software (BD Life Sciences).

### Stability of Mrf-Dendra2 signal under other conditions

*Effect of fixation on the stability of Mrf-Dendra2 signal was tested by culturing* Mtb mc^2^7000:R2:R3 in low-zinc Sauton’s medium + 100 μg/mL PAN for 7-weeks and visualizing before and after a 60-minute exposure to 4% paraformaldehyde (PFA) under microscope. *Effect of acidic pH on the stability of Mrf-Dendra2 signal* was *tested by culturing* Mtb mc^2^7000 R2:R3 in low-zinc Sauton’s medium + 100 μg/mL PAN for 7-weeks and was then transferred to acidic low-zinc Sauton’s medium + 100 μg/mL PAN (pH 4.5) for 60 minutes. Normal and acidic cells were analyzed by flow cytometry. *Effect of hypoxia on the stability of Mrf-Dendra2 signal* was *tested by culturing* Mtb mc^2^7000 R2:R3 was cultured in low-zinc Sauton’s medium + 100 μg/mL PAN for 7-weeks. Cells were then washed and placed in a tube filled to 9/10th of the total tube volume with filter sterilized supernatant of biofilm culture containing methylene blue (1.5 μg/mL) as an oxygen indicator. This Wayne model of hypoxic culture was incubated with a stir bar on a stirrer until the culture decolorized. Cells from both normal and hypoxic cultures were then analyzed via flow cytometry to assess the stability of the reporter under hypoxic conditions.

### Fluorescence-activated cell sorting

Cells of *M. smegmatis* reporter strain R2 were grown in low-zinc Sauton’s medium for 120 hours. Cell aggregates were strained and removed using sterile 5mL Falcon Polystyrene Round-Bottom Tubes with 35 μm Strainer Caps (Corning, # 352235). Cells were diluted with PBS and sorted using Becton Dickinson FACSAria II Cell Sorter. Aggregates and debris were gated out. The cells were gated into two populations, dim green and bright green. For each population, roughly 200,000 cells were collected. Serial dilutions of cells were made and ∼100 cells were plated on 7H10ADC to test for viability.

### Microfluidics for single-cell real-time growth visualization

A starter culture of *M. smegmatis* R2 reporter strain was inoculated in 7H9ADC and was grown to saturation. Cells were centrifuged for 10 minutes at 4000 rpm using a Sorvall Legend XTR centrifuge with Thermo Scientific rotor (#75003180). Cells were washed thrice with PBS and subcultured until saturating levels of Dendra2 fluorescence in 50mL of Sauton’s medium containing 1μM TPEN. Reporter expression in cells was confirmed by microscopy before setting up microfluidics. An aliquot of 100μL of cells were centrifuged and washed thrice with an equal volume of PBS with 1μM TPEN, and finally resuspended at 1:100 dilution in 100μL of PBS with 1μM TPEN. 50 μL of cells were loaded into group 8 while 50 μL of PBS was loaded into group 6 of the CellASIC ONIX plate (Millipore Sigma, #B04A-03-5PK) connected to the CellASIC ONIX2 Microfluidic System. Group 8 was primed at 13.8kPa for 15 seconds, then groups 8 and 6 were run at 27.6kPa for 15 seconds to load cells, and finally group 6 was run at 6.9kPa for 30 seconds to rinse the channel. After visually establishing an appropriate cell density in the field of view, the system was flushed for 5 minutes at 34.5kPa with PBS containing 1μM TPEN. Loaded cells were perfused for 30 minutes with PBS containing 1μM TPEN at 10kPa and then with M63 medium supplemented with 0.2% glucose and 1μM TPEN (M63 TPEN) for 36 hours. Loaded cells in microfluidics chambers were visualized using a Nikon Eclipse Ti microscope with a 1.4 N.A. 63X objective and fluorescent light source (Lumincor 200, Lumicore Inc). Images were captured using a sCMOS Edge, PCO Inc. camera. Time-lapse images were collected every 1 or 2 hours for a total of 30 hours using the Nikon NIS-Elements software to capture the images. Images were analyzed using Fiji.

The elongation rates for individual cells were determined using Fiji. A region of interest (ROI) was drawn around an individual cell to include the area with the final cell length, and the ROI was duplicated into a new window. The channels were split and the DIC channel was retained. The Yen threshold algorithm(63) was applied to binarize the ROI and assign foreground (white) and background (black) pixels with the bacterium of interest as foreground pixels. Any cells or objects in the background were subtracted so that only the bacterium of interest was visible in the ROI. The binary image was then skeletonized using “Skeletonize” function, using the following pipeline: Process: Binary: Skeletonize. The images were analyzed and measured using the “Measure_Skeleton_Length plugin(64) along with a custom-built macro “skelton length.ijm”. These measurements were and plotted in GraphPad Prism to plot the individual cellular growth rates in green and non-green cells.

### TMT-based quantification of Mtb ribosomes

Ribosomes were purified from biofilm cultures of Mtb grown in high- and low-zinc Sauton’s medium for 7-weeks. 100 μL of ribosomes at 1 μg/μl in 20mM HEPES-K pH 7.5, 30mM ammonium chloride, 10mM MgCl_2_, 5mM β-mercaptoethanol (BME) were submitted to Creative Proteomics for TMT (Tandem Mass Tags) quantification of differential ribosomal protein composition in C+ and C- ribosomes. Ribosomal proteins were reduced using 10 mM TEAB (triethylammonium bicarbonate buffer) at 56 °C for 1 hour and alkylated using 20 mM IAA (iodoacetamide) at room temperature in the dark for 1 hour. Free trypsin was added to the ribosomal protein solutions at a 1:50 ratio, and the ribosomes were digested at 37 °C overnight. The extracted proteins were lyophilized and re-dissolved in 100 mM TEAB. To label the resulting peptides, first, the TMT10plex Isobaric Label Reagent Set, Pierce Quantitative Colorimetric Peptide Assay (Thermo Fisher) was equilibrated at room temperature, 20 μL of anhydrous acetonitrile was used to reconstitute each tube within the reagent set, the reagents were dissolved for 5 minutes with occasional vortexing, and the tubes were briefly centrifuged to gather the solutions. The TMT reagents were added to each tube containing either high or low zinc derived ribosomal peptides and incubated at room temperature for 1 hour. The reactions were quenched with 8 μL of 5% hydroxylamine for 15 minutes and samples were then combined into a new microcentrifuge tube. Samples were fractionated using HPLC (high-pressure liquid chromatography). The fractionation was using Nanoflow UPLC: Ultimate 3000 nano UPLC system (ThermoFisher Scientific, USA); with a trapping column (PepMap C18, 100Å, 100 μm×2 cm, 5μm) and an analytical column (PepMap C18, 100Å, 75 μm×50 cm, 2μm). The sample volume was 5 μL. The mobile phase consisted of A: 0.1% formic acid in water and B: 0.1% formic acid in 80% acetonitrile at a total flow rate of 250 nL/min. The LC linear gradient went from 5 to 7% buffer B in 2 min, from 7% to 20% buffer B in 80 min, from 20% to 40% buffer B in 35 min, then from 40% to 90% buffer B in 4 min. Mass spectrometry (MS) was performed using a Q Exactive HF mass spectrometer (Thermo Fisher Scientific, USA) with a spray voltage of 2.2 kV and a capillary temperature 270 °C. The MS parameters were: MS resolution: 120000 at 200 m/z, and MS precursor m/z range: 200.0-1650.0. The MS/MS parameters were: Product ion scan range: start from m/z 100, Activation Type: CID, Min Signal Required: 1500.0, the Isolation Width: 3.00, the Normalized Coll. Energy: 40.0, Default Charge State: 6, Activation Q: 0.250, Activation Time: 30.00, and Data dependent MS/MS: up to top 15 most intense peptide ions from the preview scan in the Orbitrap. The MS files were analyzed using Maxquant (1.5.6.5)(65) against the *Mycobacterium tuberculosis* protein database(66). The parameters used were: the protein modifications were either carbamidomethylation (C) (fixed) or oxidation (M) (variable). The enzyme specificity was set to trypsin with the maximum missed cleavages set to 2. The precursor ion mass tolerance was set to 10 ppm, and MS/MS tolerance was set to 0.6 Da. Protein verification was performed only on highly confident identified peptides. A total of 225 proteins were identified and a quantitation ratio of low-zinc to high-zinc Mtb samples less than 1.5 was considered down-regulated while a ratio greater than 1.5 was considered up-regulated.

### Metabolomics

Starter cultures of *Mycobacterium tuberculosis* (mc^2^7000) *wt*, *Δmpy*, and *Δmpy* complement (*ΔRv_3241c harboring* pYL146) were grown planktonically in 20mL of Middlebrook 7H10 + 10% OADC + 100 μg/mL PAN + 0.05% Twee80 at 37 °C until an OD 0.4-0.6. Cells were centrifuged and washed once with detergent-free Sauton’s medium + 100 μg/mL PAN. Washed cells were resuspended in 20 mL detergent free Sauton’s medium + 100 μg/mL PAN. A 1:100 dilution of cells was inoculated detergent-free Sauton’s medium + 100 μg/mL PAN without supplemental zinc, and about 4mL was dispensed in each well of multiple 12-well plates. Pellicle biofilms of *M. tuberculosis* (mc^2^7000) and the variant strains were grown by incubating the plates for 7-weeks at 37 °C without shaking. Three independent sets of each strain were prepared. As necessary, 7-week biofilms of a subset of cultures were exposed to indicated concentration of isoniazid. Metabolite extraction was performed as previously described(67). 1 mL of freshly prepared cold metabolite extraction buffer (acetonitrile:methanol:water:: 2:2:1) was mixed with approximately 300 μL of zironica beads in a 1.5mL screw cap tubes on dry ice. Biofilms were lifted using sterile disposable 210 mm smartSpatulas^®^ (Heathrow Scientific, # 6290-120003) and flash frozen in the tube with metabolite extraction buffer and stored at -80 °C until extraction. For extraction, thawed samples were broken in a bead-beater at 6000rpm, 6 times, at 30s per round. Samples were centrifuged at 13,000 X *g* for 10 minutes at 4 °C. The supernatants from each sample were transferred to Spin-X columns and centrifuged at 5000 X *g* for 5 minutes at 4 °C and transferred to fresh sterile microcentrifuge tubes and frozen until LC-MS analysis. For LC-MS analysis, lysates were thawed on ice and crashed with one part Solvent B (100% acetonitrile containing 0.2% acetic acid). Samples were then vortexed and centrifuged at 15,000 RPM for 8 minutes at 4 °C. Supernatants were transferred to LC-MS vials and loaded onto a 4 °C temperature-controlled LC-MS autosampler. In brief, samples were separated by liquid chromatography on an Agilent 1290 Infinity LC system by injection of 2 μL of extract through a Cogent Diamond Hydride HPLC Column, 4um, 2.1 × 150 mm, 100A (Microsolv Technology Corporation). Solvent A (100% water containing 0.2% acetic acid) and Solvent B (100% acetonitrile containing 0.2% acetic acid) were infused at a flow rate of 0.400 ml min−1. The 24-min normal phase gradient was as follows: 0–2 min, 85% B; 2-3 min, 80% B; 3–5 min, 80% B; 5-6 min, 75% B; 6-7 min, 75% B; 7-8 min, 70% B; 8-9 min, 70% B; 9-10 min, 50% B; 10-11 min, 50% B; 11-11.10 min, 20% B; 11.10-14 min, 20% B; 14-14.10 min, 5% B; 14.10 min to 24 min, 5% B followed by a 10-min post-run at 85% B. Acquisition was performed on an Agilent G6546A Q-TOF mass spectrometer (Agilent Technologies) using an Agilent Jet Stream electrospray ionization source (Agilent Technologies) operated at 3500 V Cap and 1000 V nozzle voltage in extended dynamic range, in both polarities. The following settings were used for negative mode acquisition: The sample nebulizer set to 45 psi with sheath gas flow of 11 L min–1 at 350 °C. Drying gas was kept at 320 °C at 8 L min−1. Fragmentor was set to 125 V, with the skimmer set to 50 V and Octopole Vpp at 400 V. Samples were acquired in centroid mode at 1 spectra/s for *m/z* values from 50 to 1200. The following settings were used for positive mode acquisition: The sample nebulizer set to 45 psi with sheath gas flow of 11 L min–1 at 350 °C. Drying gas was kept at 320 °C at 8 L min−1. Fragmentor was set to 125 V, with the skimmer set to 50 V and Octopole Vpp at 400 V. Samples were acquired in centroid mode at 1 spectra/s for *m/z* values from 50 to 1700. Raw data was acquired from the instrument and analyzed using the Masshunter Profinder and qualitative analysis software’s. Metabolites were confirmed using a compiled database with known retention times for standards.

### Ribosome purification

Cells *of M. smegmatis* WT or the R2 reporter strain were cultured in 500ml of either low-zinc or high-zinc Sauton’s media. After 24- or 96-hours of growth, cells were harvested at 8000 rpm for 20 minutes at 4 °C in a Thermo Scientific™ Fiberlite™ F12-6 × 500 fixed-angle rotor and flash frozen in liquid nitrogen. Frozen cells were milled 6 times at 15 Hz for 3 minutes in a mixer mill (Retsch MM400). The milled cells were resuspended in 15 mL of cold HMA-10 buffer (20 mM HEPES-K pH 7.5, 30 mM NH4 Cl, 10 mM MgCl2, 5 mM β-mercaptoethanol) and centrifuged at 15,000rpm for 30 minutes at 4 °C in a Thermo Scientific F21-8 x 50y fixed angle rotor. The supernatants were transferred into the Beckman PC ultracentrifuge tubes (Beckman 355618) and centrifuged at 42,800rpm for 2 hours and 15 minutes at 4 °C in a Ty70Ti, Type 70 Ti Fixed-Angle Titanium Rotor (339722) Beckman rotor. The supernatants were discarded, and the resulting pellets were soaked in 1mL of HMA-10 overnight. The soaked pellets were resuspended in the 2 mL HMA-10 buffer. Pellets were then homogenized for 20 minutes on ice and transferred to low speed 40 mL centrifuge tubes. Homogenates were diluted with another 2 mL of HMA-10 and treated with 3 units/mL TURBO™ DNase (2 U/μL) (Invitrogen™, #AM2238) for one hour at 4 °C. To the DNase treated homogenates, 4mL of HMA-0.06 buffer (20 mM HEPES-K pH 7.5, 600 mM NH4 Cl, 10 mM MgCl_2_, 5 mM β-mercaptoethanol) and incubated at 4 °C for 2 hours to remove peripheral proteins. The homogenates were then centrifuged at 13,000 rpm for 15 minutes at 4 °C. The supernatants were transferred into the Beckman PC ultracentrifuge tubes and centrifuged at 42,800 rpm for 2 hours and 15 minutes at 4 °C in a Ty70Ti rotor. The supernatants were discarded, and the resulting pellets containing the crude ribosomes were soaked in 500 μL of HMA-10 buffer overnight. The pellets were resuspended in the 1 mL HMA-10 buffer. To purify the ribosomes, the samples were centrifuged at 13,000 rpm for 15 minutes at 4 °C in a benchtop centrifuge (Eppendorf 5415D). The ribosomes were quantified by measuring absorbance at 260 nm. For separating individual subunits, up to 600 μL of ribosomes were layered on top of a 40 mL 10% to 40% sucrose gradient in HMA-10 buffer and centrifuged for 16 hours at 24,000 rpm in a Beckman rotor SW28. The overnight gradients were fractionated using the Brandel density gradient fractionation system to separate the individual subunits. Fractions corresponding to the 70S ribosomes were pooled separately, diluted with HMA-10, and pelleted by ultracentrifugation at 42,800 rpm for 4 hours at 4 °C in a Beckman rotor Type 70Ti rotor. The pellets were soaked overnight in 500μL HMA-10 buffer and diluted with another 500μL HMA-10 buffer for a final volume of 1mL of purified 70S ribosomes. The 70S subunit was quantified by measuring the 260 nm absorbance and stored at -80 °C until further use.

### Mpy Immunoblotting

2.4 picomoles of 70S ribosomes from indicated strains were mixed with 4X SDS loading buffer for a total volume of 1 mL and heated for 10 minutes at 95 °C, resolved in 15% SDS-PAGE gel and were transferred to PVDF membranes using the Bio-Rad Trans-Blot SD Semi-Dry Transfer Cell for 70 minutes at 105 mA. The membranes were blocked with 5% nonfat milk overnight, washed, and probed for 1 hour with indicated antibodies: S13 (DSHB, mouse, 5 μg/mL corresponding to 1:100); S14C- (in-house, rabbit, 1:2000); Mpy (in-house, rabbit, 1:5000) in 1% nonfat milk. The membranes were washed and then incubated with goat anti-rabbit secondary HRP conjugated antibody (1:5000) or anti-mouse secondary HRP conjugated antibody (1:5000). The membranes were developed with ECL reagents (Thermo Scientific, #PI32106) and exposed to chemiluminescence films, which were developed and scanned for visualization.

### RADA staining and visualization

#### Staining cells from batch culture

*M. smegmatis WT, Δmpy,* or *Δmpy*-complemented cells expressing R2 reporter were grown in 50mL low-zinc media for approximately 96 hours. After growth, samples were inoculated at an OD of 0.5 in 7H9 + 0.5% glycerol + 10% ADC + 0.05% Tween-80 and incubated at 37 °C on a shaker for 15 minutes to initiate outgrowth. After 15 minutes, 50 μM RADA (5-carboxytetramethylrhodamine amino-D-alanine; Tocris, #6649), was added and incubated at 37 °C on shaker for an additional 15 minutes. Cells were then centrifuged in a benchtop centrifuge (Eppendorf 5425) for 2 minutes at 13,000rpm and washed thrice with an equal volume of PBST. After the final wash, cells were resuspended in PBS without tween. Coverslips were treated with 1mg/mL Poly-D-lysine hydrobromide, 2.5 μL of culture was applied to the treated coverslip, and slides were mounted with SlowFade Diamond (Thermo Fisher, #S36963). Cells were visualized immediately using a Nikon Ti2-E w/iLas2 coupled to a Lumecor SPECTRA Light Engine and the accompanying NIS Elements Advanced Research version 6.10.01software. Images were captured using Photometrics Prime BSI Express sCMOS 4.2-megapixel camera. Five fields were acquired per sample. Cellpose(68) was used for cellular segmentation and to define the regions of interest (ROI) for each bacterium using the acquired phase images. Each ROI was manually verified, and any incorrect ROIs were edited or deleted, while any omitted ROIs were manually added using FIJI. A Python script was written to determine the mean green fluorescence, green or non-green category, red mean fluorescence, red max fluorescence, polar red mean fluorescence, polar red max fluorescence, polar enrichment, localization, and classification of the cells as either green or non-green with either polar or diffuse red signal. The cutoff for defining a cell as P1 (green) or P2 (non-green) was determined using the background signal exhibited by a high-zinc culture. Polar enrichment was defined as a signal 1.3 times higher at the pole than the rest of the bacterial body. Fields were deconvolved using Autoquant (Media Cybernetics) using the Gibson-Lanni theoretical point-spread-function (PSF) and then analyzed using Fiji. For Mtb, mc^2^7000 and *Δmpy* strains expressing R2 reporter were grown in low-zinc Saution’s containing supplemented with 100 μg/mL PAN. A high-zinc sample was used to determine the background autofluorescence cutoff. After two weeks, 1 μM TPEN was added to each culture, and cultures were grown for another two weeks. After two weeks, a portion of the media from each culture was filter-sterilized, and cells were resuspended at an O.D. of 0.5 in 1 mL of filter-sterilized media containing 50 μg/mL isoniazid and incubated at 37 °C for one week. Cells were then washed and resuspended in 500 μL of 7H9 + 10%OADC + 0.5% glycerol + 0.05% Tween-80 + 100 μg/mL PAN containing 50 mM RADA overnight. After incubation, cells were centrifuged in a benchtop centrifuge (Eppendorf 5425) for 2 minutes at 13,000 rpm and washed thrice with an equal volume of PBST. After the final wash, cells were fixed with 4% paraformaldehyde for 1 hour, centrifuged and resuspended in PBS and mounted as described for *M. smegmatis*. Cells were visualized immediately using a Dragonfly 200 Series spinning disk confocal with accompanying Fusion software. Images were captured using a Zyla sCMOS camera. Three to five fields were acquired per sample. Samples were deconvolved using Autoquant (Media Cybernetics) using the Gibson-Lanni theoretical point-spread-function (PSF). Fiji was used to determine the polar mean fluorescent intensities of RADA staining in mycobacterial strains to assess differences in growth potential. To determine polar staining, the pixel intensities were measured along the length of each bacillus using the Plot Profile function. A Python script was written to collate the individually measured values along the bacterial rods across three to five fields. The polar values were defined as the pixel values within the first micron of the bacterium. The average pixel intensities were calculated for each bacterium’s pole and plotted.

#### Staining cells in microfluidics

*M. smegmatis* cells were prepared and loaded in a CellASIC ONIX plate (Millipore Sigma, #B04A-03-5PK) and connected to the CellASIC ONIX2 Microfluidic System using the procedure described above. The loaded cells were perfused for 30 minutes with PBS containing 1μM TPEN at 10kPa. M63 medium supplemented with 0.2% glucose and 1μM TPEN (M63 TPEN) and containing 50μM RADA was perfused for 1 hour, washed with PBS-TPEN for 30 minutes, and visualized. Second and third cycles of 2- and 4-hour RADA exposure, washing and visualization were similarly performed on the same cells. Microfluidics were visualized using a Nikon Eclipse Ti microscope with 1.4 N.A. 63X objective and fluorescent light source (Lumincor 200, Lumicore Inc). Images were captured using a sCMOS Edge, PCO Inc. camera. Time-lapse was recorded using the Nikon NIS-Elements software to capture an image every 15 minutes for 7 hours. Polar RADA staining in individual cells was quantitively analyzed using Fiji software using the “Plot Profile” function to measure pixel intensities as described above.

### qRT-PCR

Total RNA was extracted from *M. smegmatis* cultures using TRIzol (Invitrogen, #15596026). Genomic DNA was depleted with two sequential rounds using the TURBO DNA-free™ Kit (Invitrogen™, #AM1907). The first strand cDNA was synthesized using the Maxima cDNA synthesis kit (Thermo Scientific, #FERK1641). Real time PCR was performed on an ABI 7000 instrument with SYBR Green master mix as per the manufacturer’s instructions. Briefly, 20ng of cDNA, 1μL of 5μM primers were mixed with 10μL of the master mix in a total volume of 20μL, and the PCR was performed at: 95C for 10 minutes, followed by 40 cycles of 95C for 10 seconds and 60C for 1 minute. For each set of experiment, an endogenous control with *sigA*-specific primers, as well as no-template and no reverse transcriptase controls were set-up for normalization.

### Antibiotic susceptibility *in vitro*

#### Survival of under isoniazid

*M. tuberculosis mc^2^7000 wt* and *Δmpy* expressing R2 were grown in low-zinc Sauton’s medium for 7 weeks at 37 °C. After 7 weeks, wells were pooled and washed three times with PBS + 0.1% tyloxapol. Cells were dispersed overnight using 3mm glass beads at 4 °C in PBS + 0.05% tyloxapol. Cells were then filtered through 40 μm cell strainers (CellTreat, #229481) to remove large aggregates, yielding homogenous suspension. The OD of each sample was determined, and cells were resuspended in 12mL PBS + 0.05% tyloxapol with or without supplemental zinc. Isoniazid (50 μg/mL) or streptomycin (5 μg/mL) and rifampicin (50 μg/mL) was added to cultures. Cells were incubated at 37 °C and once a week, cells were serial diluted and plated to enumerate colonies.

#### Growth inhibition by isoniazid

##### Isoniazid minimum inhibitory concentration 90 (MIC 90) of mc^2^7000, Δmpy, and complemented strains

Mtb mc^2^7000, isogenic *Δmpy,* and complemented strains were grown in 7H9 + 10% OADC + 0.5% glycerol + 0.05% Tween-80 + 100 μg/mL PAN until an O.D. between 0.3 and 0.5. Cells were then washed and resuspended at a 1:10 dilution in Sauton’s media + 100 μg/mL PAN with serial diluted concentrations of isoniazid and grown for 1 week at 37 °C. The O.D. was measured at each concentration, and the precent growth was determined against the no-antibiotic control. The MIC 90 was calculated using the Lambert and Pearson method in which the proportion of bacteria at each concentration was plotted against the log-transformed antibiotic concentration and fit to a Gompertz nonlinear regression model to determine the MIC (69).

### RNA-Seq

Total RNA was extracted from *M. tuberculosis* cultures using RNAeasy Mini Kit (QIAGEN Inc., #74104). Genomic DNA was removed using the TURBO DNA-free™ Kit (Invitrogen™, #AM1907). RNA integrity was determined using a Qubit 2.0 Fluorometer and a Tapestation 2200. Ribosomal RNA was depleted using TruSeq Stranded Total RNA with Illumina Ribo-Zero Plus rRNA depletion (Illumina, #20040526). Libraries were prepared using NEBNext Ultra II Directional RNA Library Prep Kit for Illumina (NEB, #E7765). Adaptors were ligated used NEBNext Multiplex Oligos for Illumina (Index Primers Set 3 (NEB, #E7710S). Libraries were sequence on an Illumina NextSeq 500/550 and sequences were analyzed using Rockhopper(70).

### Animal infection, antibiotic treatment, necropsy and lung tissue analysis

All animal experiments were conducted in accordance with the protocols approved by the Institutional Animal Care and Use Committee of the Wadsworth Center (#21-447) and Albany Medical College (#21-03002 and #21-04003). C3HeB/FeJ and C57BL/6 mice were infected with *Mtb* (Erd):R2:R3 or isogneic *Δmpy*:R2:R3 expressing Mrf-Dendra2 from pYL200 and mCherry from plasmids pEM2(19) or pJL016, respectively. Infections were performed in the ABSL3 laboratory at Wadsworth Center using a Glas-Col aerosolizer (Glas-Col Inc.) with settings (10 minutes nebulization, 15 minutes of cloud time and 10 minutes of exhaust) that deposited ∼100-200 bacilli in each mouse(19). After 12- and 14-weeks of infection for C3HeB/FeJ and C57BL/6, respectively, mice were euthanized and lung sections were processed for immunofluorescence staining, immunophenotyping, microscopic visualization of R2 reporter expression in bacteria, and determining total bacterial burden using the methods described below.

#### Bacterial burden

For enumerating bacterial burden in the lungs, one of the lung lobes was homogenized in PBS + 0.05% Tween-80, strained through a 40 μm cell strainer, and the homogenates were serial diluted and plated on 7H11OADC agar. As necessary, 50 μg/mL hygromycin or 20 μg/mL kanamycin were added in the plate. Colonies were counted after 4 weeks of incubation at 37 °C.

#### Microscopic analysis of bacilli expressing R2 reporter

A portion of the lungs was fixed in 0.4% paraformaldehyde in PBS (Fisher Scientific) at room temperature for 7 days. Fixed lungs were soaked in 15% sucrose in PBS overnight at 4 °C and then soaked in 30% sucrose in PBS overnight at 4 °C. Tissues were mounted into blocks using Scigen Tissue-Plus O.C.T. Compound (Fisher HealthCare) while submerging the blocks in a slurry of dry ice in ethanol. Frozen tissue blocks were cryosectioned at 12 μm thickness on a CryoStarNX70 (Thermo Scientific). Before microscopy, sections were thawed for 5 minutes at room temperature and washed with PBS for 10 minutes in coplin jars, thrice, to remove O.C.T media. Sections were mounted overnight in the dark with Molecular Probes™ ProLong™ Diamond Antifade Mountant (Thermo Fisher, #P36961) to allow the mounting agent sufficient time to polymerize. Mounted sections were visualized by Leica SP5 confocal laser scanning microscope with a 60X oil immersion objective. Dendra2 was excited with a 488 nm argon laser and mCherry was excited with a 594 nm laser. Dendra2 was detected by photo multiplier tube (PMT) at 496-550 nm while mCherry was detected using a HyD detector at 600-650 nm range. Images were captured with Leica SP5 camera using Leica Microsystems Confocal LAS AF software. Images were deconvolved using Autoquant (Media Cybernetics) using the Gibson-Lanni theoretical point-spread-function (PSF) and then analyzed using Fiji. Mrf-Dendra2 signal predominantly appears as foci. Rigorous colocalization analysis of mCherry signal with Mrf-Dendra2 was performed to ensure that observed green signals were originating from Mtb cells. For colocalization, an optical slice with a green signal was chosen from the z-stack, a line was drawn through the green signal and RGB Profiler was used to determine if the green signal colocalized with the constitutively labeled red bacilli. For signals to be colocalized, the difference between distance of the red and the green signal peaks must be no more than one pixel. The colocalized red and green signal was used as a criterion for designating Mrf-Dendra2 positive bacterium. Depending on bacterial load, 4 to 9 independent fields of view were analyzed per mouse. From each necropsy timepoint 2 male and 2 female mice were imaged.

#### Immunofluorescence staining with anti-Ly6G antibody and DAPI

Frozen OCT sections were removed, thawed, and washed 4 times with PHEM + 0.5% Tween80 for 5 minutes each. The sections were permeabilized with PHEM + 0.5% Triton X-100 for 10 minutes, washed 3 times for 5 minutes each with PHEM + 0.5% Tween80, and blocked PHEM + 5% rat serum + 5% w/v BSA + 0.1% Triton X-100 for 1 hour at room temperature. Permeabilized and blocked sections were soaked in IgG2a, κ rat anti-mouse Ly6G Clone 1A8 conjugated to Alexa-647 (BioLegend, #127609) diluted 1:500 in PHEM + 5% rat serum + 5% w/v BSA + 0.1% Triton X-100 overnight at 4 °C. The stained sections were then washed three times with PHEM + 0.5% Tween80 and mounted using ProLong Diamond with DAPI (Thermo Fisher, #P36962) overnight at room temperature. The mounted sections were visualized on a Dragonfly 200 Series spinning disk confocal microscope with accompanying Fusion software. Images were captured using a Zyla sCMOS camera, deconvolved using Autoquant (Media Cybernetics) using the Gibson-Lanni theoretical point-spread-function (PSF) and analyzed using Fiji.

#### Immunophenotyping

Mouse lungs were transferred to a small petri dish containing 5 mL PBS and were washed twice with 3 ml of fresh PBS. The lungs were then diced in an Eppendorf tube containing 1 ml PBS. To each mixture, a pre-warmed digestion cocktail was added and the mixture was incubated at 37 °C for 1 hour on rocker. The digests were passed through a 40 µm cell strainer and the resulting liquids were centrifuged at 500 x g for 5 minutes at 4 °C. If the cell pellets looked bloody, then 1 mL of RBC lysis (ACK lysis) buffer was added. The cells were incubated on ice for 3 minutes, 10 mL of 5% FBS in RPMI were then added, the suspensions were centrifuged at 500 x g for 5 minutes, and the supernatants were discarded. The cells were then washed twice with 1 mL FACS buffer by centrifuging at 500 x g for 5 minutes at 4 °C. A cell count was performed and 1 million cells from each mouse were used for antibody staining. The supernatants were aspirated and the pellets were resuspended in 50 µl of Fc-block solution (1:200). The samples were incubated in dark for 20 minutes at 4 °C and then topped with 150 µL of FACS buffer. The cells were then washed by spinning at 500 x g for 5 minutes at 4 °C. The supernatants were aspirated, and the pellets were resuspended in 50 µL of staining buffer (CD cocktail) containing fluorescence conjugated antibodies (1:100). The samples were incubated in dark for 60 minutes at 4°C then topped with 100 µL of FACS buffer and centrifuged at 500 x g for 5 minutes at 4°C. Cells were washed thrice with 800 µL FACS buffer by centrifuging at 500 x g for 5 minutes at 4°C. Cells were then fixed in 100 µL of BD Cytofix, and were incubated in dark for 30 minutes at 4 °C, after which 100 µL of FACS buffer was added, and the content was centrifuged at 500 x g for 5 minutes at 4 °C. After discarding the supernatants, the pellets were resuspended in 200 µL of BD cytofix, diluted 1:4 with FACS buffer. The samples were transferred out of BSL-3 facility and stored at 4 °C until analysis by flow cytometery. Flow Cytometry was performed using a BD FACSymphony™ A3 Cell Analyzer with the following lasers: Blue laser 488-100 LS, Red laser 637-140 LX, Violet laser 405-100 LX, and Yellow-green laser 561-100 LS.

#### Isoniazid (INH) treatment and analysis of reporter expression

Two groups of C3HeB/FeJ mice each were infected for 8-weeks with Mtb(Erd):R2:R3 or *Δmpy*:R2:R3 reporter expressing Mrf-Dendra2 from pYL200 and mCherry from plasmids pEM2(19) or pJL016, respectively. While one group of mice for each bacterial strain was treated with 100 μg/mL isoniazid (in drinking water) for four weeks, the other group was left untreated for entire four weeks. Four mice (2 male + 2 female) from the treated group were euthanized at 0-, 2-, and 4-week post INH treatment, and 4 mice from the untreated group were euthanized after 12 weeks post-infection. Homogenates from a lung lobe of a mouse were plated for determining total bacterial burden. Another portion of the lung was fixed for one-week in 4% paraformaldehyde in PBS and OCT sections were processed and visualized using a Leica SP5 confocal laser scanning microscope as described in the methods above.

#### RNA-Seq of lungs tissues and neutrophils

For transcriptomics of total lung tissues from uninfected and infected mice, a portion of lungs was collected into RNAlater (Sigma-Aldrich, #R0901) immediately after necropsy and frozen in -80 °C until RNA extraction. The tissues were disrupted using a pestle and the lysates were homogenized in lysis buffer (RLT Plus + 1% beta mercaptoethanol). The homogenates were passed through 70 µm cell strainers to remove any debris and the clear lysates were centrifuged for 10 minutes at 13,500 rpm in a benchtop centrifuge. The supernatants were removed, transferred to gDNA eliminator spin columns, and placed into a 2 mL collection tubes. The samples were centrifuged for 3 minutes at 13,500 rpm and the flow-throughs were saved and the columns were discarded. The RNAs were extracted using the Qiagen RNeasy Plus Mini Kit (Qiagen, #74134) following manufacturer’s instructions. One volumed of 70% ethanol was added to each of the flow-through and mixed by pipetting up and down. The samples, including any precipitates, were transferred to spin columns, placed into a 2 mL collection tubes, and centrifuged for 2 minutes at 13,500 rpm. The flow-throughs were discarded, and the spin columns were washed by adding 700 μL of RW1 buffer to the columns and centrifuging for 2 minutes at 13,500 rpm. The flow-throughs were discarded, and the spin columns were washed twice with 500 μL of RPE buffer for 2 minutes at 13,500 rpm. The columns were dried at room temperature and the RNA was eluted into fresh collection tubes using 50 μL of RNase-free water. The RNA concentrations were determined using a Quibit 2.0 Fluorometer and the integrities were assessed using an Agilent TapeStation 4200.

About 2 μg of total RNA from each sample was used for sequencing performed by Azenta Life Sciences. RNA sequencing libraries were prepared using the NEBNext Ultra II RNA Library Prep Kit for Illumina (NEB, Ipswitch, MA, USA) according to the manufacturer’s instructions. The mRNAs were enriched using Oligod(T) beads, then fragmented for 15 minutes at 94 °C, followed by first strand and second strand cDNA synthesis. The cDNA fragments were end-repaired and underwent 3’ adenylation followed by ligation with universal adaptors, index addition, and library enrichment using PCR. The libraries were quantified using a Quibit 2.0 Fluorometer and qPCR (KAPA Biosystems, Wilmington, MA, USA) and validated using an Agilent TapeStation 4200.

The libraries were multiplexed and clustered according to manufacturer’s instructions on the Illumina NovaSeq flowcell. The libraries were sequenced using a 2X150bp Paired End (PE) configuration. NovaSeq Control Software (NCS) was used for base calling. Raw .bcl sequence data from the lllumina NovaSeq were converted to fastq files and de-multiplexed using Illumina bcl2fastq 2.0 software with one mismatch allowed for each index sequence identification. Sequence reads were trimmed to remove adaptor sequences and poor-quality reads. The trimmed reads were mapped to the reference genome on ENSEMBL using the STAR aligner v.2.5.2b to generate BAM files. Gene hits exclusively from exons were calculated using the Counts from the Subread package v.1.5.2. Genes with counts less than 10 in all samples were discarded. DESeq2 was used to compare gene expression between groups and samples. Log2 fold changes and p-values were determined using the Wald test. Differentially expressed genes were denoted as those with adjusted p-values < 0.05 and absolute log2 fold change >1. GeneSCF was used to determine gene ontology.

For transcriptomics of neutrophils, single cell suspensions of Mtb (HN878) infected mouse lungs from C3H/FeBJ and C57BL/6 mice were harvested in cold PBS in a petri dish, on ice. Harvested lung lobes were digested at 37 ℃ in 4 mL of PBS containing Collagenase Type IV (150 U/mL) (Gibco, #17104019) and DNase I (60 U/mL) (Roche-Sigma Aldrich, #10104159001). Homogenates were strained through a 70 μM cell strainer and treated with 3 mL ammonium–chloride–potassium (ACK) to remove any remaining red blood cells. The suspensions were gently washed, and the pellets were resuspended in PBS + 0.5% BSA and blocked with 50 μL (1:400) Fc-Block CD16/32 (Bio Legend, #156604) to prevent non-specific binding, followed by treatment with APC anti-Ly6G antibody at 1:200 (BioLegend, #127613) for 30 min at 4°C in the dark. Ly6G-bound cells were then treated with 1:10 anti-APC nanobeads according to the MojoSort™Human anti-APC nanobead kit protocol (BioLegend, #480090) and stained cells were washed and collected on MojoSort™ Magnet (BioLegend, #480019). Unbound cells were discarded, and bound cells were collected in PBS + 0.5% BSA. Purity of enriched cells was determined by staining with Pacific Blue™ anti-mouse Ly6G/Ly6C GR1 antibody at 1:200 (BioLeagend, #108429) and analyzed by flow cytometry as described earlier. Purified Ly6G+ cells (totaling between 10^6^-10^7^) were stored at −80 °C in RLT Plus Buffer (Qiagen, #1053393) for RNA isolation. The RNA was isolated from these cells according to the Qiagen RNeasy Plus Mini Kit (Qiagen, #74134) protocol. RNA concentrations were determined using a Quibit 2.0 Fluorometer and integrities were assessed using an Agilent TapeStation 4200.

Between 750 ng to 1 μg οf total RNA was used for sequencing. The Qiagen Fast Select rRNA HMR Kit (Qiagen, #334385) was used to remove rRNA. Libraries were prepared using the NEBNext Ultra II RNA Library Prep Kit for Illumina (NEB, #E7770). After adaptor ligation, library integrity as assessed using a Tapestation 4200 and quantified using a Quibit 2.0. Libraries were sequenced on an Illumina HiSeq using 2X150 Pair-End sequencing.

To analyze sequences, the raw .bcl sequence files were first converted to fastq files and demultiplexed using bcl2fastq. The adaptor sequences and poor-quality reads were trimmed using Trimmomatic v.0.36 and the reads were mapped to the reference *Mus musculus* genome on ENSEMBLE using Rsubread v1.5.3. Entrez Gene IDs was used to quantify the gene counts. To determine differential expression, limma-voom was utilized. To perform this, STAR aligner v.2.5.2b mapped the genes to the ENSEMBL reference genome. Unique exon reads were selected for differential expression analysis using DESeq2 to compare the reads between the mice strains. The Wald Test generated the Log2 fold changes and their respective P-values. Gene ontology analysis was used to identify roles of genes. Genes with less than 10 reads were excluded. Specifically, genes related to zinc and metal transport, such as S100a8/S100a9 (calprotectin), and Slc30a (ZnT) and the Slc39a (ZIP) genes were examined.

#### Visualization and confirmation of extracellular bacteria

The neutrophil reporter mice used in this study were created in the Mishra Lab by breeding the B6.129(Cg)-*Gt(ROSA)26Sor^tm4(ACTB-tdTomato,-EGFP)Luo^*/J (**BL6^mT/mG^**) and B6. Cg-Tg (S100A8-cre,-EGFP)1Ilw/J (**S100a8-cre**) animals in specific pathogen free conditions at the animal resource facility of Albany Medical College (AMC) in accordance with protocols approved by the AMC Institutional Animal Care and Use Committee (IACUC) (Animal Care User Protocol Numbers: ACUP-24-03003 and 21-04003). The BL6^mT/mG^ strain contained a two-color fluorescent reporter construct that targets the cellular membranes throughout mouse tissues and labels cells with tdTomato, expressing widespread strong red fluorescence. Flanking the membrane-targeted tdTomato (mT) cassette are *loxP* sites. The S100A8 promoter was chosen as a neutrophil specific marker. When bred with S100a8-cre, the Cre recombinase expressed by the S100A8 promoter excised mT cassette in S100A8 cells, yielding specific green fluorescence through the expression of the membrane-targeted EGFP (mG) in these cells. Mice genotypes were confirmed using PCR and gel electrophoresis for site-specific alleles. Flow cytometry using a using a BD FACSymphony™ A3 Cell Analyzer was used to confirm green fluorescence in neutrophils (CD11b^hi^/Ly6G^hi^ positive cells).

Chimeric mice were infected with Mtb HN878 expressing pAO621. Infections were performed in the ABSL3 laboratory at Albany Medical College using a Glas-Col aerosolizer (Glas-Col Inc.). After 42 days of infection, mice were euthanized and lung sections were processed for immunofluorescence staining, immunophenotyping, microscopic visualization as described above. For microscopic visualization, sections were visualized on a Dragonfly 200 Series spinning disk confocal microscope and Fusion software. Images were captured using a Zyla sCMOS camera, deconvolved using Autoquant (Media Cybernetics) using the Gibson-Lanni theoretical point-spread-function (PSF), and processed using Fiji. Visualized green fluorescence was further validated to be originating from neutrophils by performing immunofluorescent staining with IgG2a, κ rat anti-mouse Ly6G Clone 1A8 conjugated to Alexa-647 (BioLegend, #127609) as described above. The HN878 BFP reporter infection lacks a constitutive fluorescent reporter. Visible blue fluorescence reports the subset of bacteria that have undergone ribosome remodeling. The distance between an individual bacillus and its nearest neutrophil and non-neutrophil was determined in FIJI, and the distances were plotted in GraphPad.

#### Ribosome remodeling in S100a9^-/-^ mice

Five C57BL/6 and five C57BL/6: s100a9^-/-^ mice (∼10 week of age) were infected with a reporter strain of Mtb(Erd):R1:R3, harboring a plasmid pYL28 that expressed Dendra2 from the Zur-regulated promoter of C- ribosome (Mtb:pEM2:pYL28) and plasmid pEM2 that expressed mCherry from the constitutive *hsp60* promoter. Infection was dosed at ∼100 CFU/mouse using the Glas-Cole aerosolizer using the default settings described for the R2 reporter. At 14-weeks post-infection, four males and one female of C57BL/6 and three males and two females of 5 C57BL/6: s100a9^-/-^ were euthanized and lungs were processed for determining bacterial burden and for confocal microscopy as described for the R2 reporter.

#### Zinc starvation response in Mtb in neutropenic mouse model

Six male and six female C3HeB/FeJ mice (9-10week of age) were infected with Mtb(Erd):R1:R3. Infection was dosed at approximately 450 CFUs/mouse using a Glas-Col Inhalation Exposure System. At 1-week post-infection, 6 mice (3 males & 3 females) were administered with an anti-mouse Ly6G (Clone 1A8, BioX-Cell, USA), while the remaining 6 mice received an isotype control (rat IgG2a BioX-Cell, USA). All antibodies were administered through intraperitoneal injection once daily at 200µg/mouse for 15 days. One day after the final dose, mice were euthanized and lungs were processed for gross bacterial burden, immunophenotyping, and confocal microscopy as described for the R2 reporter strain.

### Statistical analysis

With the exception of RNA-Seq experiments, all statistical tests and analyses were performed using GraphPad Prism Versions 9 and 10 (Dotmatics). To assess differential RADA staining in P1 and P2 populations in WT *Δmpy,* and/or *Δmpy::comp*, unpaired Kruskal-Wallis ANOVA tests were performed with N=5 independent fields for *M. smegmatis* and N=3 for *M. tuberculosis*. Densitometric scan of immunoblots showing differential puromycin incorporation was statistically analyzed using a paired two-tailed t-test with N=5 biological replicates. For determining P-values for comparisons of Mrf-Dendra2:mCherry or Dendra2:mCherry ratios, and the CFUs per lung in mouse studies (N=4-6 mice per group), a two-tailed Mann-Whitney test was used. For determining the P-values for comparing the distance between bacteria and neutrophils or non-neutrophils in the chimeric mouse model, a two-tailed Mann-Whitney test was used. Depending on bacterial burden, 7-9 fields were analyzed per mouse. Differences in MSH/MSSM were assessed using paired t-tests with three independent biological replicates. For assessment of differential antibiotic sensitivity in Mtb wild-type and *Δmpy* strains, unpaired t-tests were used for two independent biological replicates. For RNA-seq datasets, q-values between two conditions (N=3) were determined using the default Wald test in DESeq2.

## Acknowledgments

Authors acknowledge technical help from Dr. Fioranna Renda for image analysis. This work was supported by grants to AKO (NIH: AI132422, NIH: AI163599), BBM (NIH: R01HL166257), and KYR (NIH: AIR25140472, NIH: AIP01159402). JHC was partially supported by the RNA Fellowship of the RNA Institute, University at Albany. JH was supported by APHL through a Cooperative Agreement Number NU60OE000104 (CFDA #93.322), funded by the Centers for Disease Control and Prevention (CDC) of the US Department of Health and Human Services (HHS). Breeders for *S100a9^-/-^*strain, which was originally created by Thomas Vogl at the Institute of Immunology in Münster, Germany, was donated by Thomas Kehl-Fie of UIUC, Urbana-Champaign, IL, USA. Service and support are also acknowledged from the Wadsworth Center’s Applied Genomic Technologies Cluster, Advanced Light Microscopy, Immunology, and Media and Tissue Culture Core facilities.

## Authors’ Contributions

JHC, JH, VP, YL, PS, BBM and AKO designed and performed biochemical, genetics and animal experiments; JHC, AS, AW and KYR performed metabolomics experiments; JHC, JH, RWC and AKO acquired and analyzed micrographs. All authors wrote the manuscript.

## Conflict of Interest

Authors declare no conflict of interest.

## Data availability

The authors declare that data supporting the findings of this study are available within the paper. Raw sequencing datasets for the RNA-seq experiments for Mtb, neutrophils, and mouse lung homogenates are deposited at Gene Expression Omnibus (GEO) with accession number GSE263900, GSE244230 and GSE263940, respectively. The reviewer token for accessing the datasets are khspieqybfovvol, etmbgigmtdaxhyx, and ybslcyoybpmrhad, respectively.

## Supplementary Materials

**Figure S1:**
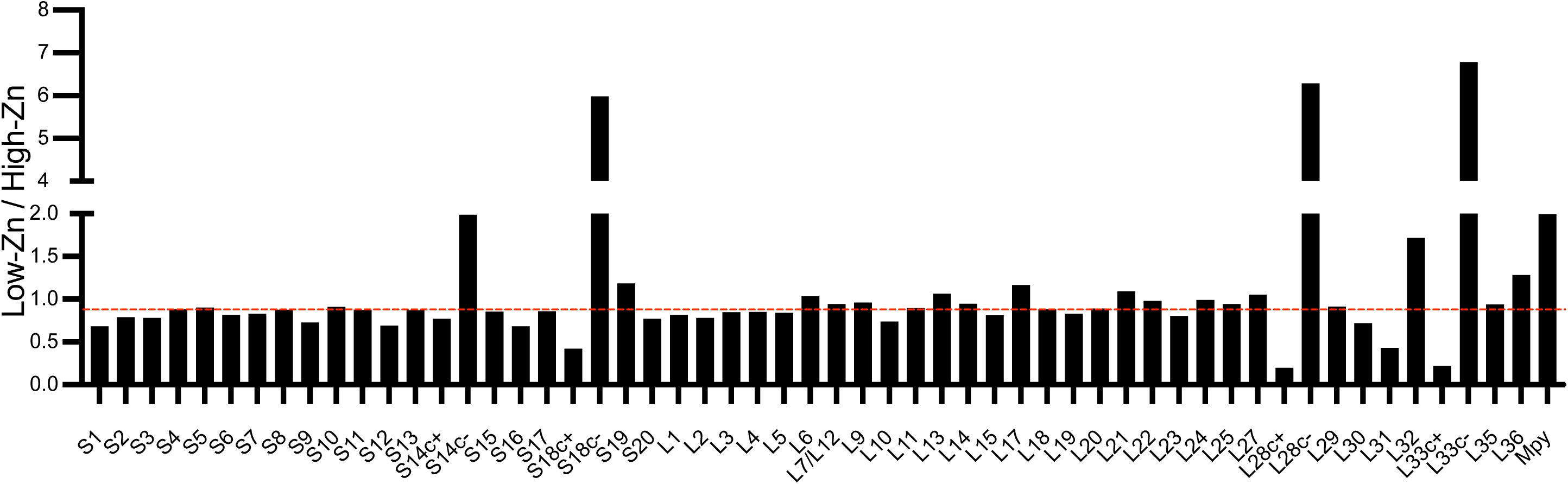
Ratio of ribosomal proteins and Mpy in the 70S ribosomes from Mtb (mc^2^7000) cultured for 7-weeks as biofilms in low- and high-zinc Sauton’s medium (denoted as Low-Zn and High-Zn, respectively). Relative levels of proteins in the two ribosomes were determined by tandem mass tag mass spectrometry (TMT-MS). Red line indicates the median ratio of all invariant ribosomal proteins.

**Figure S2:**
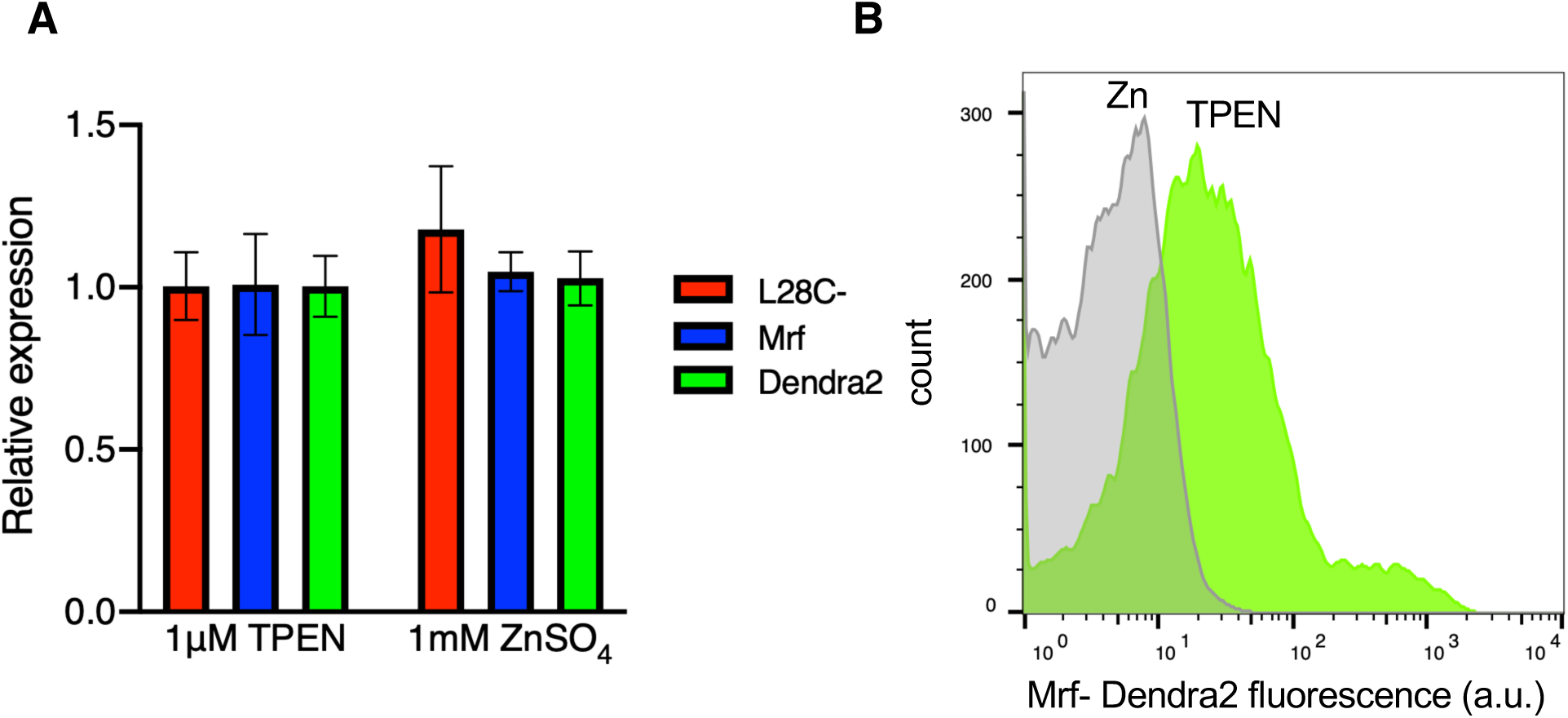
Post-transcriptional regulation of Mrf-Dendra2 in R2 reporter of *M. smegmatis*. **A.** Relative levels of Mrf and Dendra2 transcripts expressed from the corresponding regions in *P_mrf_-*Mrf-Dendra2 (R2) reporter in a *Δzur* strain cultured in high-zinc (1 mM ZnSO_4_) and low-zinc (1μM TPEN) conditions for 96 hours. SigA was used as endogenous invariant control for normalization. **B.** Analysis of Mrf-Dendra2 signal by flow cytometry in high- and low-zinc cultures of the R2 reporter of *Δzur: P_mrf_-*Mrf-Dendra2 cells after 96 hours of growth.

**Figure S3:**
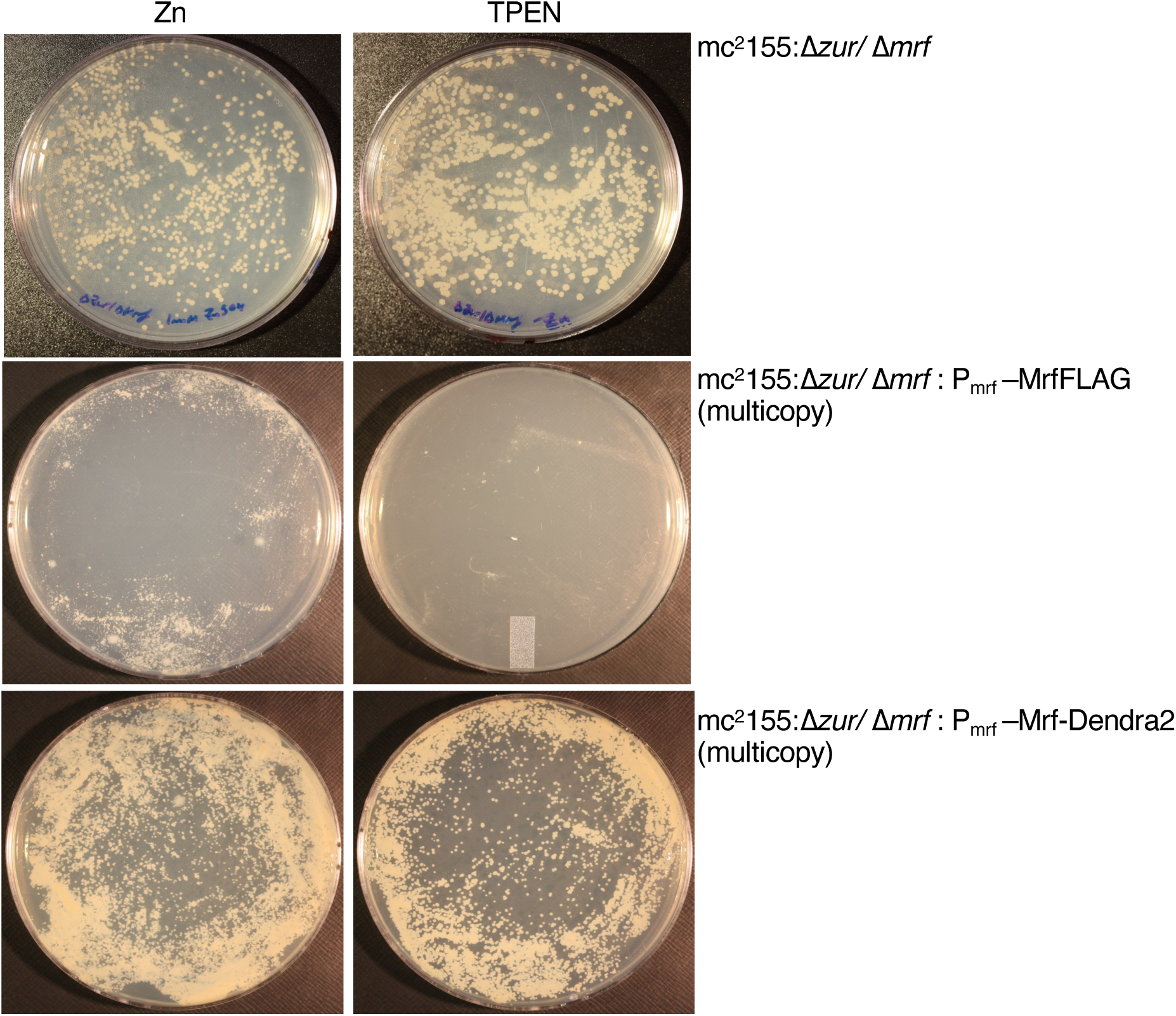
Loss of Mrf function in Mrf-Dendra2. Plating efficiency of *M. smegmatis* mc^2^155: Δ*zur/*Δ*mrf* upon transformation with a multicopy plasmid constitutively expressing either Mrf-FLAG (pYL141) or Mrf-Dendra2 (pYL167) on a high- or low-zinc Sauton’s medium, marked as Zn and TPEN, respectively. Unlike Mrf-FLAG, which is toxic at high levels in zinc-starved cells, Mrf-Dendra2 is well tolerated.

**Figure S4:**
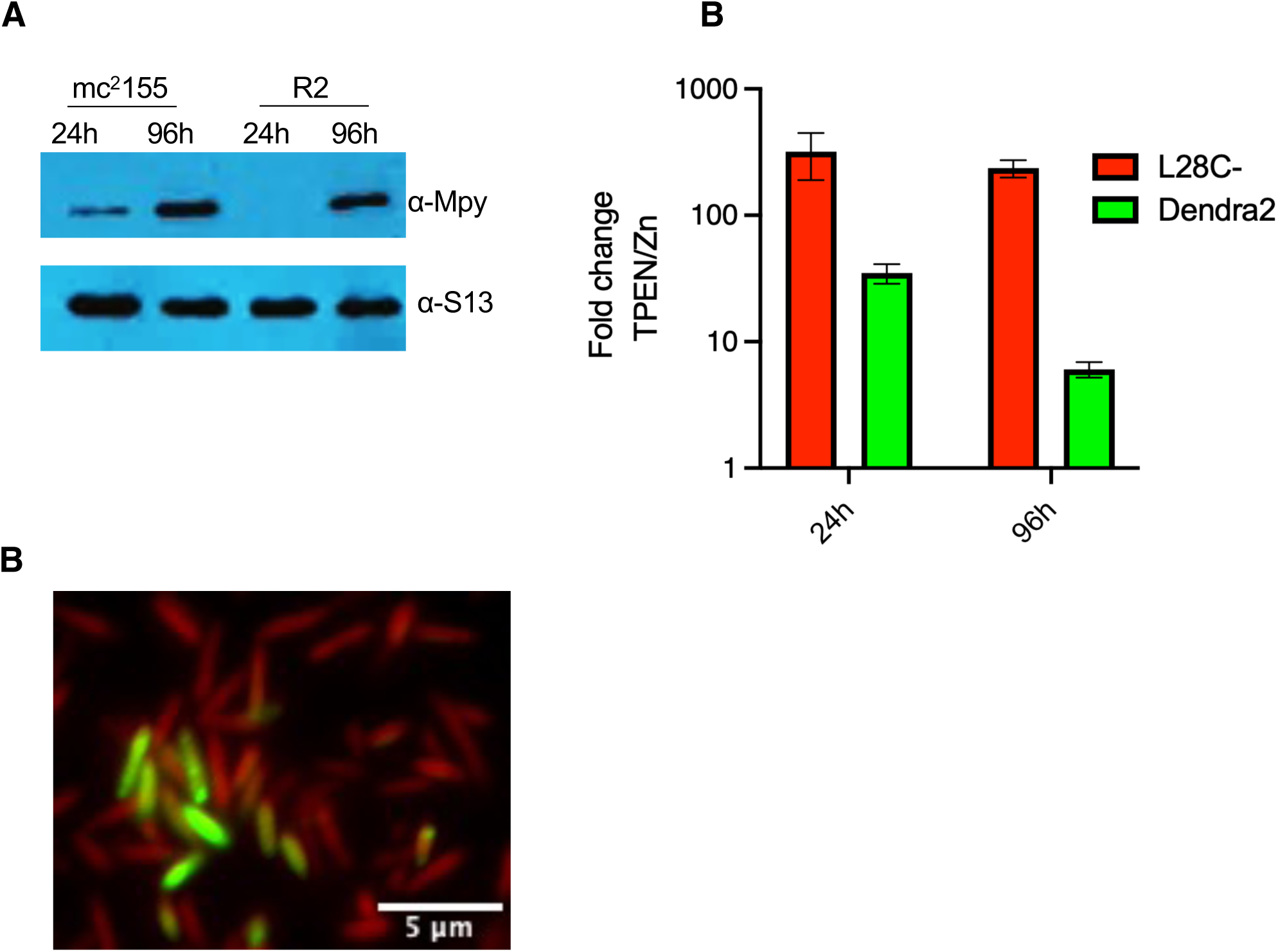
Concurrence of Mrf-Dendra2 accumulation and Mpy recruitment to the ribosome in *M. smegmatis*. **A.** Mpy levels in the ribosome purified from the parent *M. smegmatis* mc^2^155 cells and its R2 reporter after 24- and 96-hours of growth in low-zinc Sauton’s medium. The ribosomal protein S13 was probed to normalize total ribosomes in each well. **B.** Dendra2 transcript levels in the R2 reporter at 24- and 96-hours of growth in low-zinc Sauton’s medium, relative to the level in high-zinc (1mM ZnSO_4_) Sauton’s medium. L28c-transcripts were measured as a reference for C-operon. SigA was used as endogenous invariant control for normalization. **C.** Representative micrograph of bacilli with Mrf-Dendra2 signal in *M. smegmatis*.

**Figure S5:**
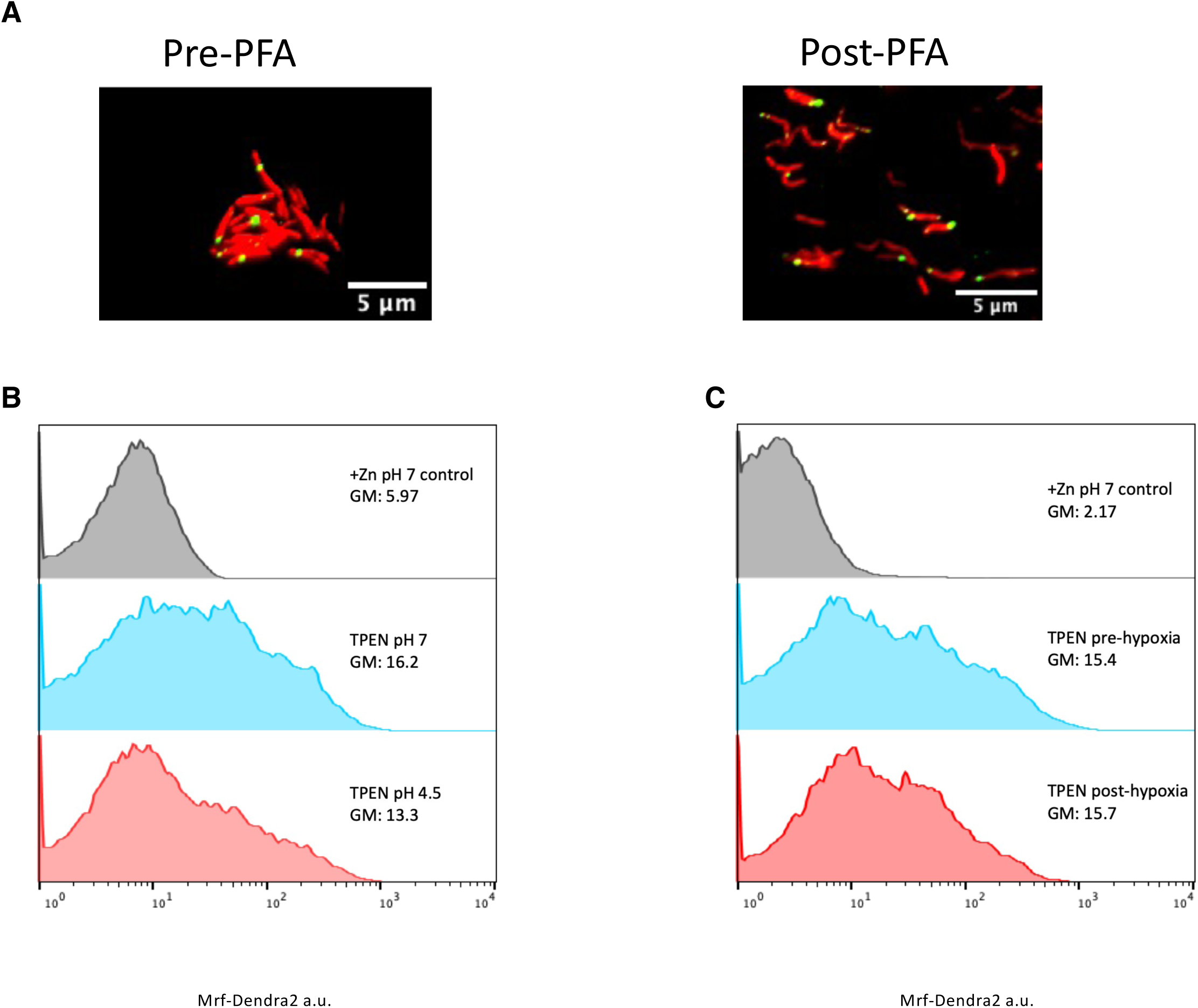
Stability of Mrf-Dendra2 in zinc-starved Mtb mc27000 after exposure to other conditions. **A.** Effect of fixation on the stability of Mrf-Dendra2 signal. Micrographs of the mc^2^7000:R2:R3 reporter cultured in low-zinc Sauton’s medium for 7-weeks and visualized before and after a 60-minute exposure to 4% paraformaldehyde (PFA). **B.** Effect of acidic pH on the stability of Mrf-Dendra2 signal. R2 reporter of mc^2^7000 cultured in low-zinc Sauton’s medium for 7-weeks was then transferred to acidic low-zinc Sauton’s media (pH 4.5) for 60 minutes, after which both normal and acidic cells were analyzed by flow cytometry. **C.** Effect of hypoxia on the stability of Mrf-Dendra2 signal. An aliquot of R2 reporter of mc^2^7000 cultured in low-zinc Sauton’s medium for 7-weeks was transferred to a tube to fill up to 9/10^th^ of the total tube volume and methylene blue (1.5 μg/mL) was added as an oxygen indicator. This Wayne model of hypoxic culture was incubated on a stirrer until the cultured decolorized. Cells from both normal and hypoxic cultures were then analyzed via flow cytometry to assess the stability of the reporter under hypoxic conditions. Cells from high-zinc cultures were used as negative control to set the base line for flow cytometry in panel B and C. GM denotes geometric mean of Mrf-Dendra2 signal.

**Figure S6:**
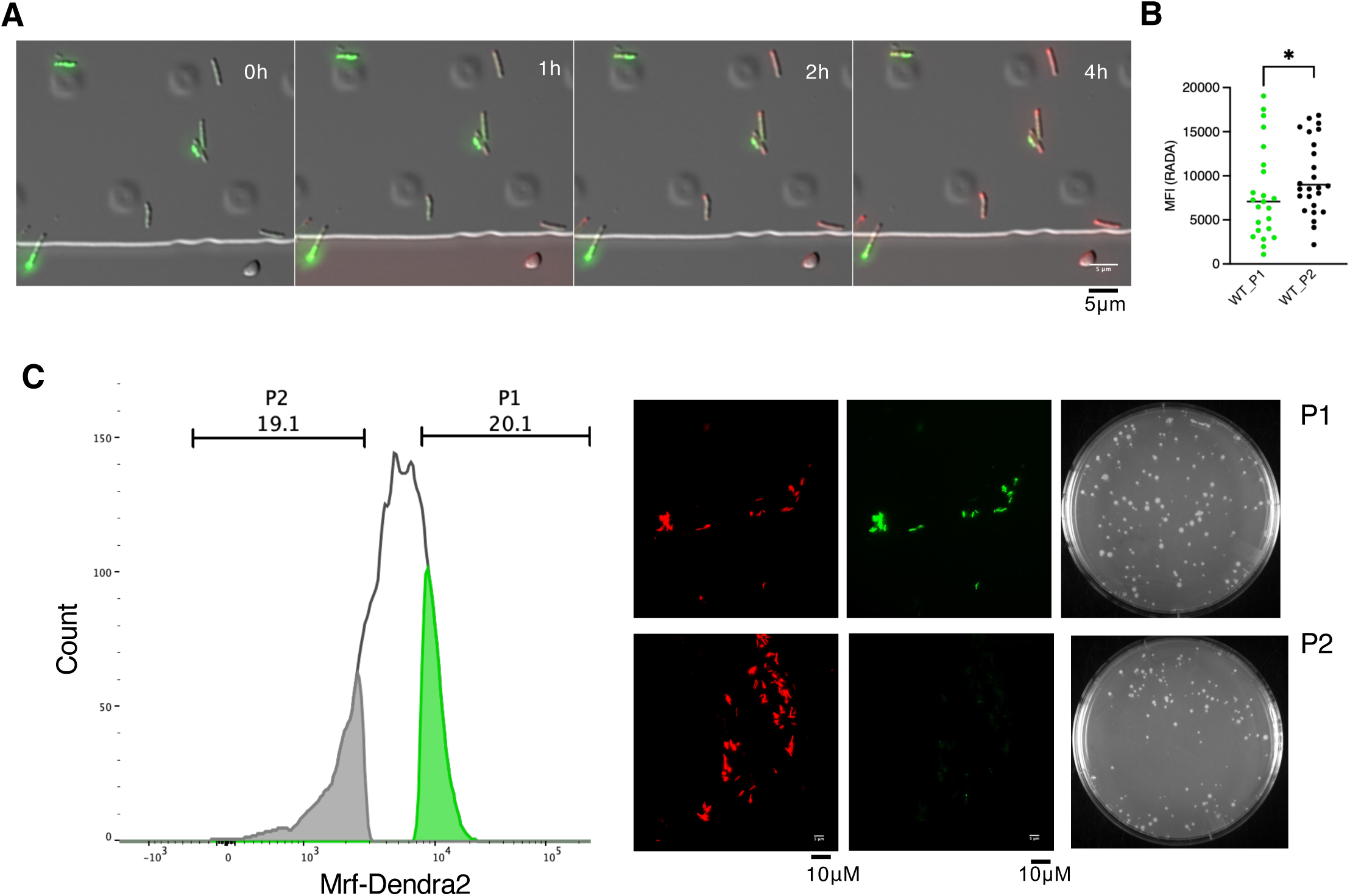
Mrf-Dednra2 accumulation and metabolic slowdown in viable *M. smegmatis* cells. **A-B.** Snapshots from a time-lapse visualization of RADA staining in P1 and P2 cells of R2 reporter of *M. smegmatis* mc^2^155 in a microfluidic chamber. Cells from 120-hour-old batch culture in low-zinc Sauton’s medium were loaded and PBS was perfused for 30 minutes prior to perfusing M63 minimal medium containing RADA for 1-, 2- and 4-hours, washed and visualized. Quantitative analysis of the mean fluorescence intensity (MFI) in individual P1 and P2 cells after 2 hours of RADA exposure. * indicates P < 0.05 (Mann-Whitney). **C.** Validation of the viability in P1 and P2 cells by their plating efficiency. 20 % of the cells with the most and least Mrf-Dendra2 fluorescence intensity (based on the histogram shown on the left) were sorted. After confirming the reporter signals under microscope, ∼100 cells were plated on 7H10ADC to enumerate the colonies.

**Supplementary Movie 1A (attached separately):**
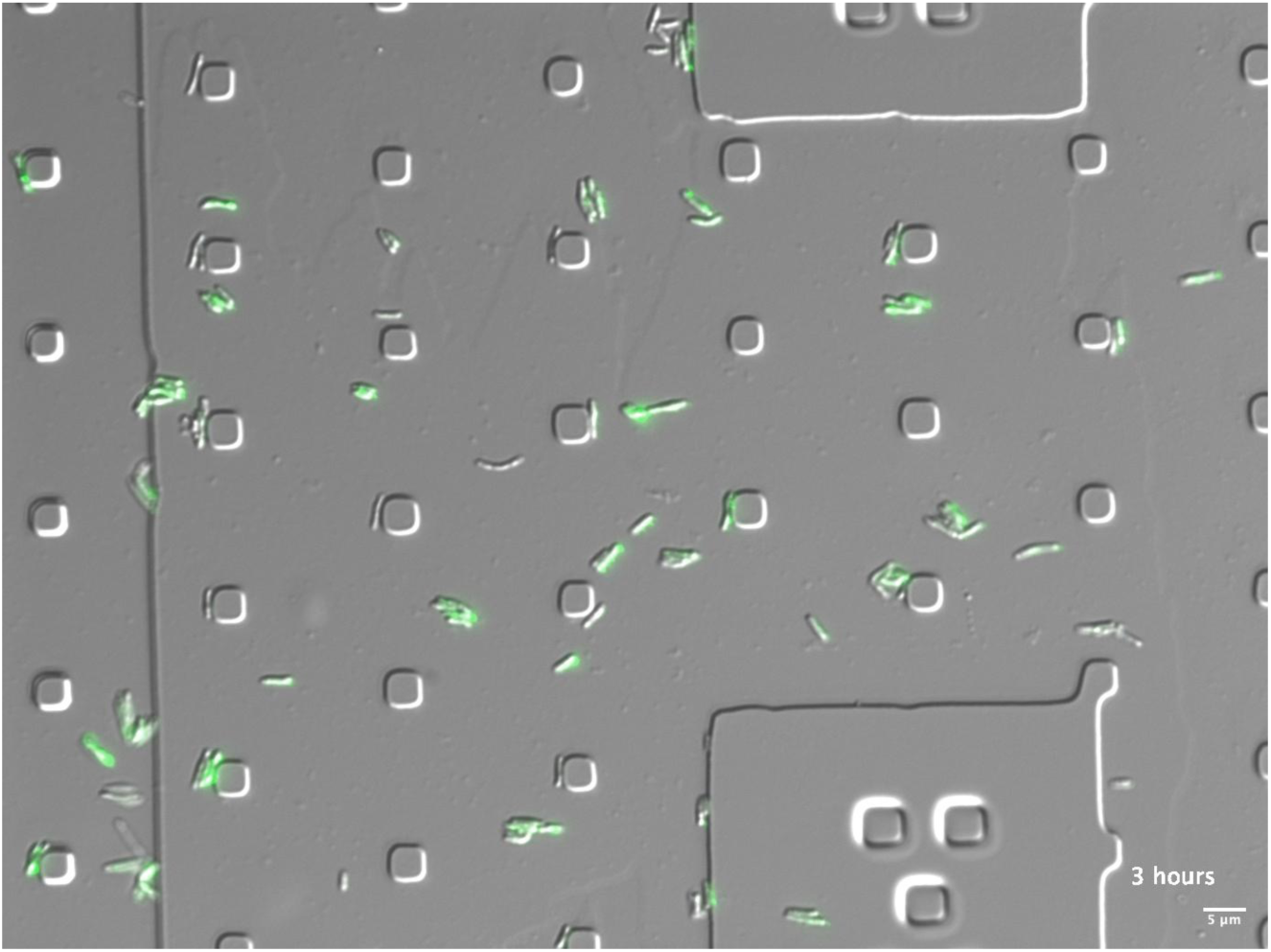
A time-lapse observation of the growth of P1 (green) and P2 (non-green) cells of the R2 reporter of mc^2^155 strain of *M. smegmatis* in a microfluidic chamber. Cells obtained from 120-hour low-zinc cultures of the two strains were washed and perfused in the chamber with M63 minimal medium containing 1 µM TPEN. This placeholder shows just the first frame of the entire movie, while the movie is uploaded separately as a supplemental file.

**Supplementary Movie 1B (attached separately):**
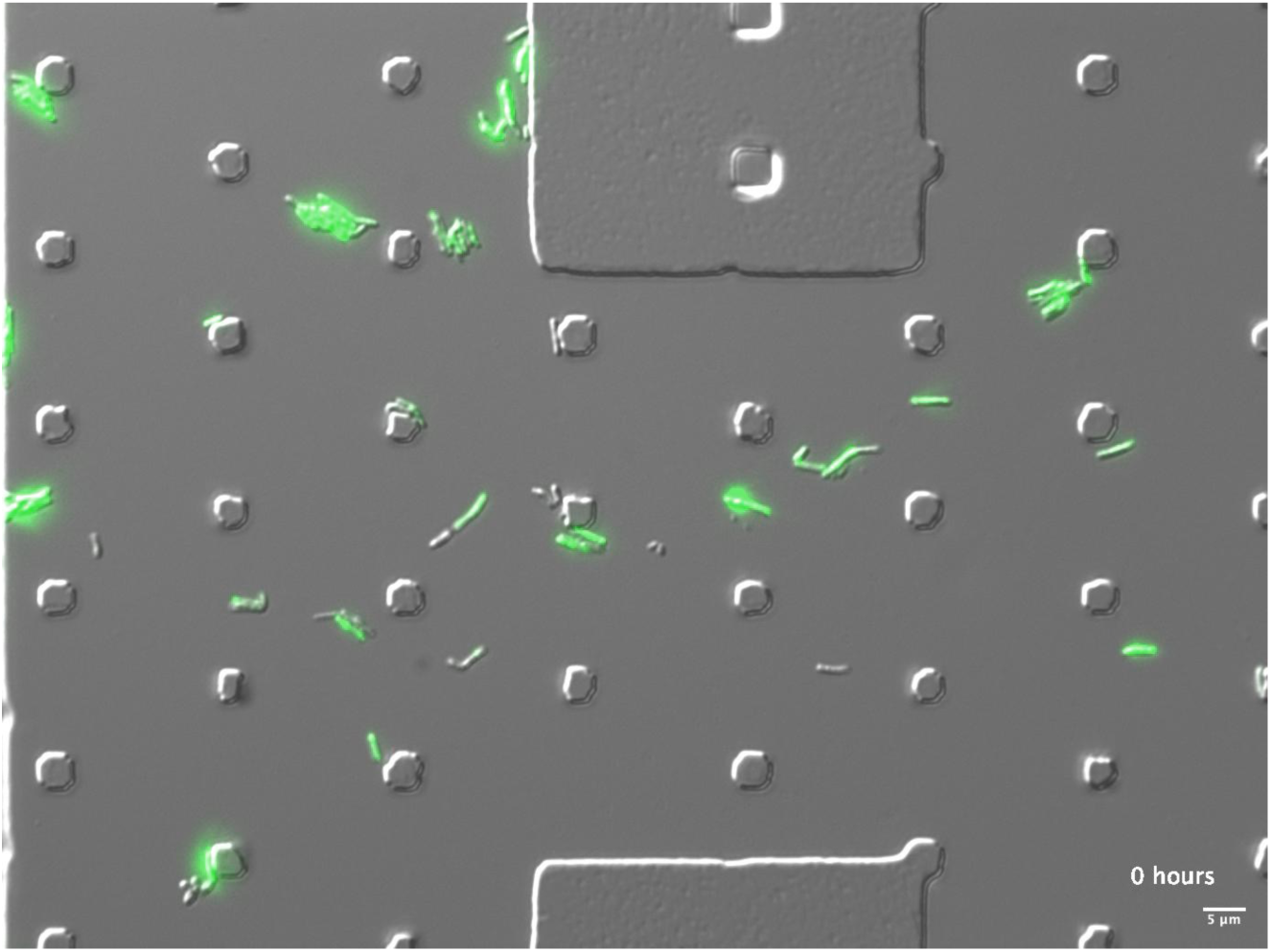
A time-lapse observation of the growth of P1 (green) and P2 (non-green) cells of the R2 reporter of *Δmpy* strain of *M. smegmatis* in a microfluidic chamber. Cells obtained from 120-hour low-zinc cultures of the two strains were washed and perfused in the chamber with M63 minimal medium containing 1 µM TPEN. This placeholder shows just the first frame of the entire movie, while the movie is uploaded separately as a supplemental file.

**Figure S7:**
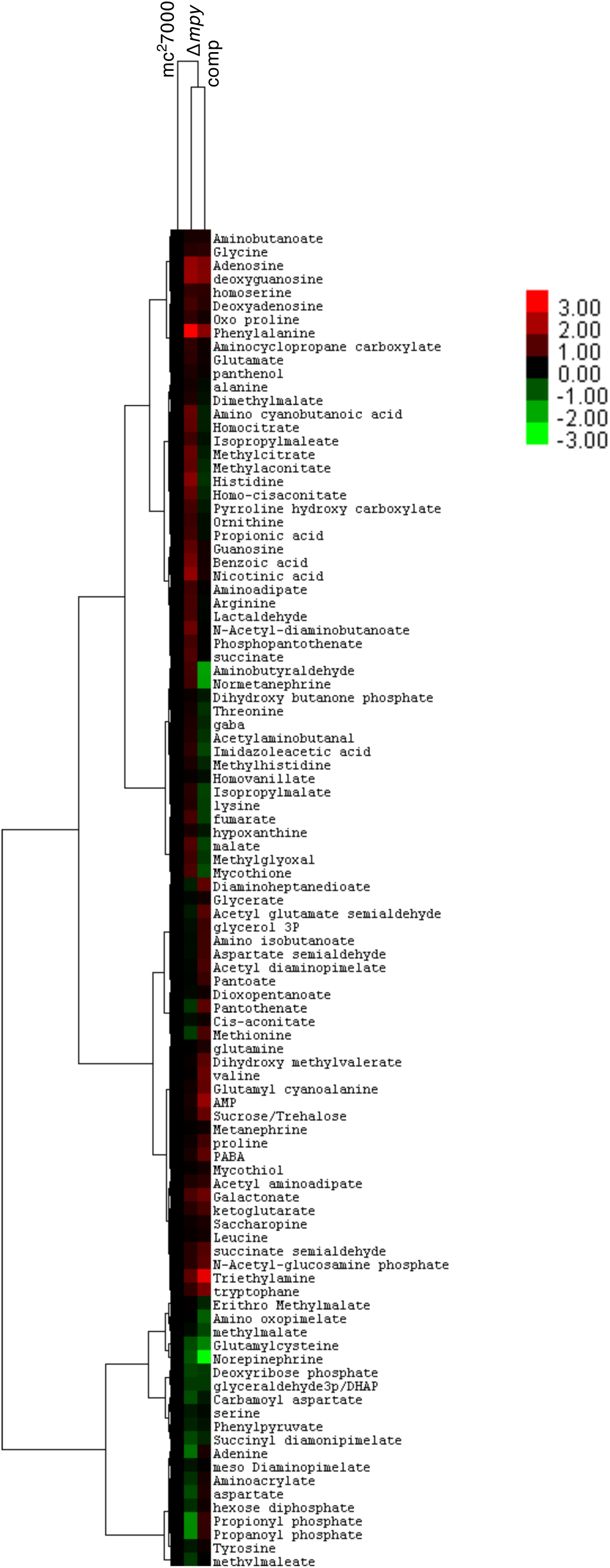
Profile of differentially abundant metabolites from Mtb mc^2^7000, isogenic *Δmpy,* and complemented (comp) strains from 7-week biofilm cultures.

**Figure S8:**
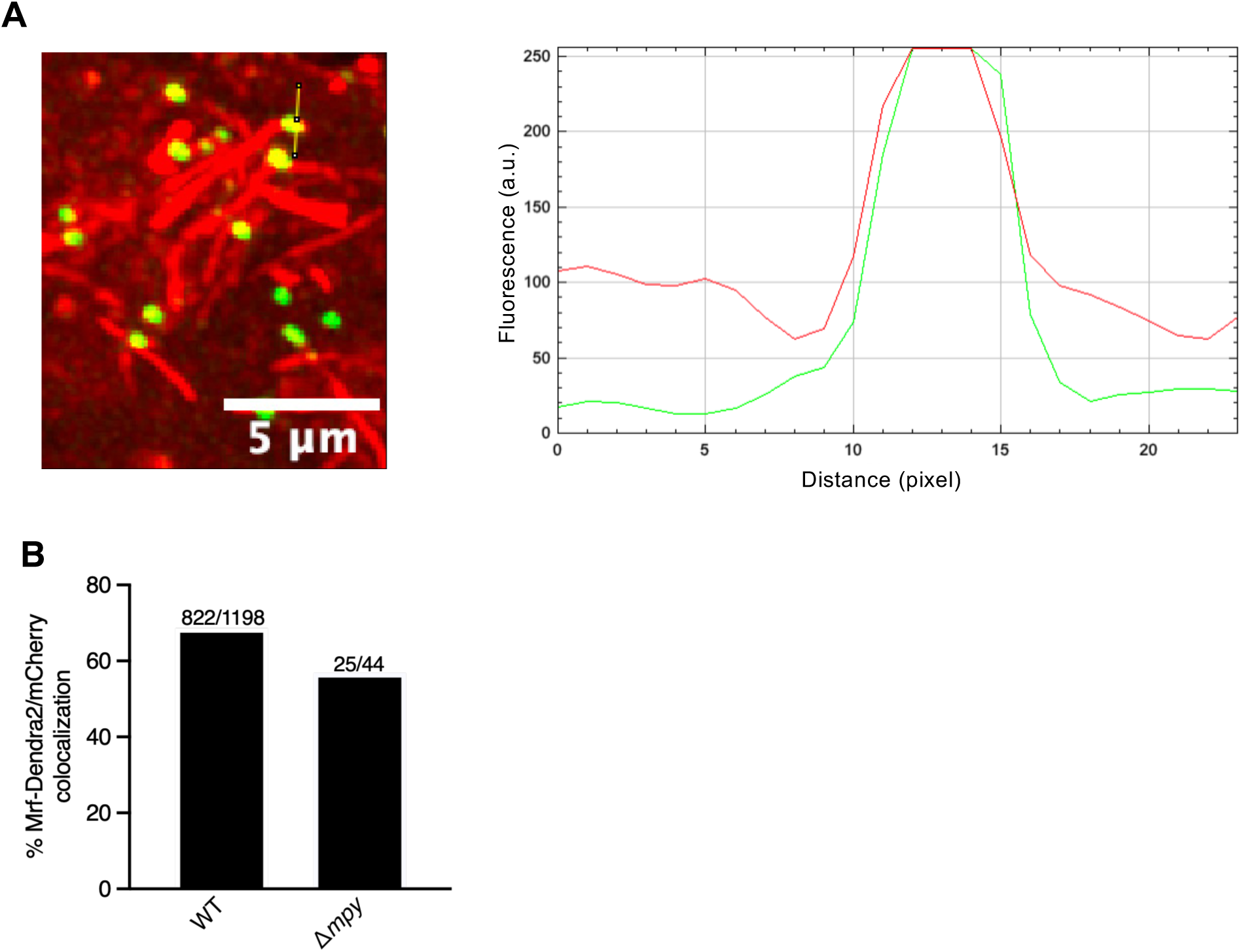
Colocalization analysis of red and green fluorescent signal from mCherry and Mrf-Dendra2, respectively, in Mtb(Erd):R2:R3 reporter in lung sections described in figure 3. **A.** RGB plot profile of red and green fluorescence in a representative image shown on the left. **B.** Average frequency of colocalized mCherry and Mrf-Dendra2 signal obtained from multiple sections from multiple infections with Mtb(Erd):R2:R3 and *Δmpy*:R2:R3 reporter strains in C3HeBFeJ mouse strain. The numbers at the top indicate the total number of bacilli analyzed.

**Figure S9:**
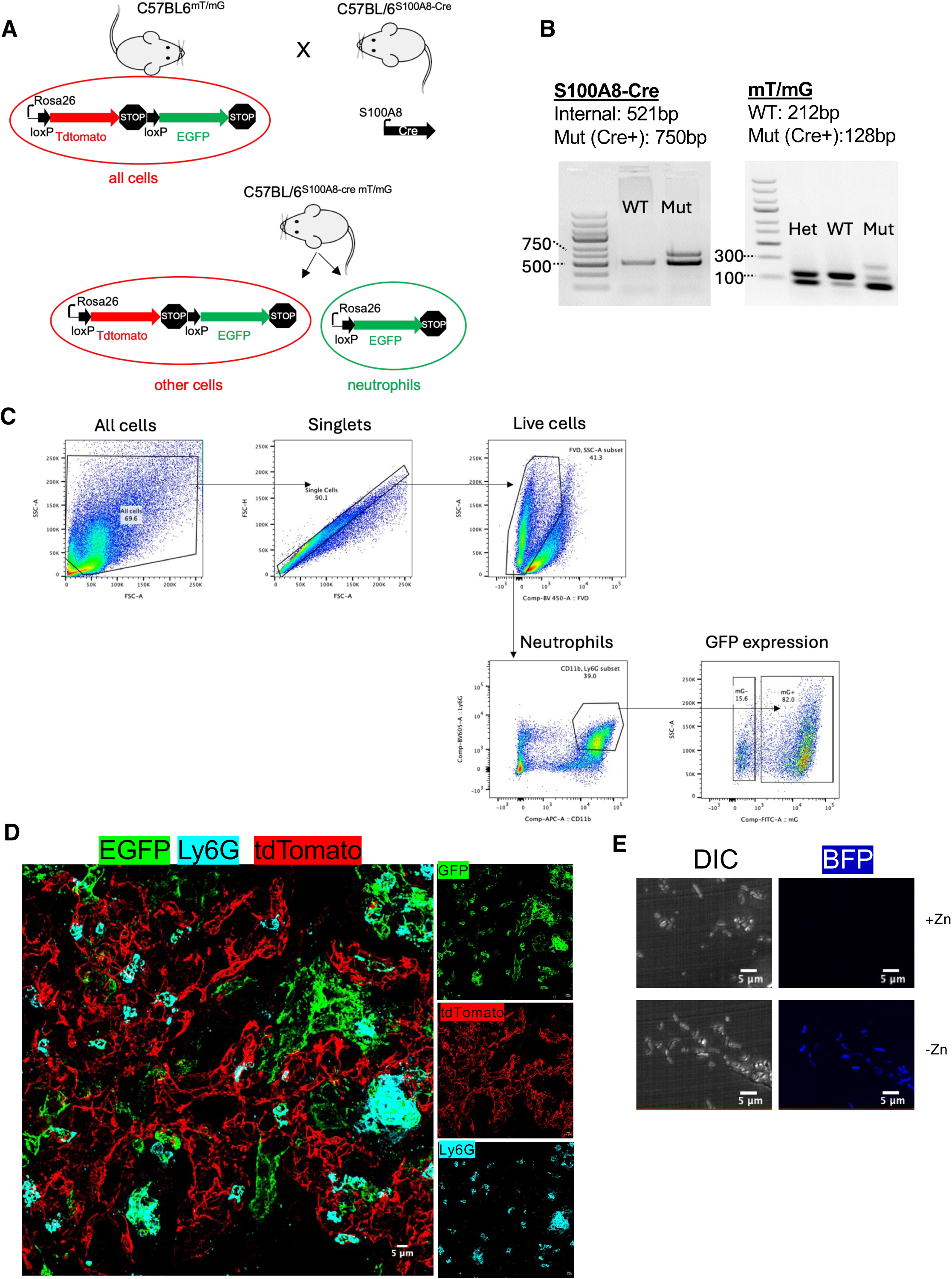
Generation of neutrophil reporter mice. **A.** A breeding scheme between BL6*^mT/mG^* and *S100a8-cre* generating the neutrophil reporter mice (S100a8cre^mT/mG^) where Cre expressing cells specifically labeled with membrane tagged GFP while other cells remained tdTomato positive. **B.** PCR-based genotyping of the S100a8-cre^mT/mG^ showing the Cre positivity and homozygous mT/mG allele. The respective wild-type (WT) parents were used as controls. A heterozygous strain (Het) for mTmG is also shown. **C.** Confirmation of GFP expression in lung neutrophils (Ly6G+ cells) obtained from S100a8-cre^mT/mG^ mice by flow cytometry. **D.** Co-localization of neutrophil-specific marker, Ly6G, (visualized with Alexa Fluor 647-conjugated anti Ly6G antibody staining, cyan) mGFP cells (green) in a Mtb-infected lung sections of S100a8-cre^mT/mG^ mouse. Other cells in the lung are labeled with tdTomato (mT). Scale bar, 5 µm. **E.** Zinc-responsive expression *in vitro* of blue fluorescent protein (BFP) in a Mtb(HN878):R4 reporter strain carrying a transcriptional fusion of BFP from C-r-protein promoter. The reporter strain was grown on Sauton’s agar plate with or without supplemental zinc (6 µM ZnSO4) and imaged under bright (DIC) and UV light (BFP; ex – 374 nm, em – 443 nm).

**Figure S10:**
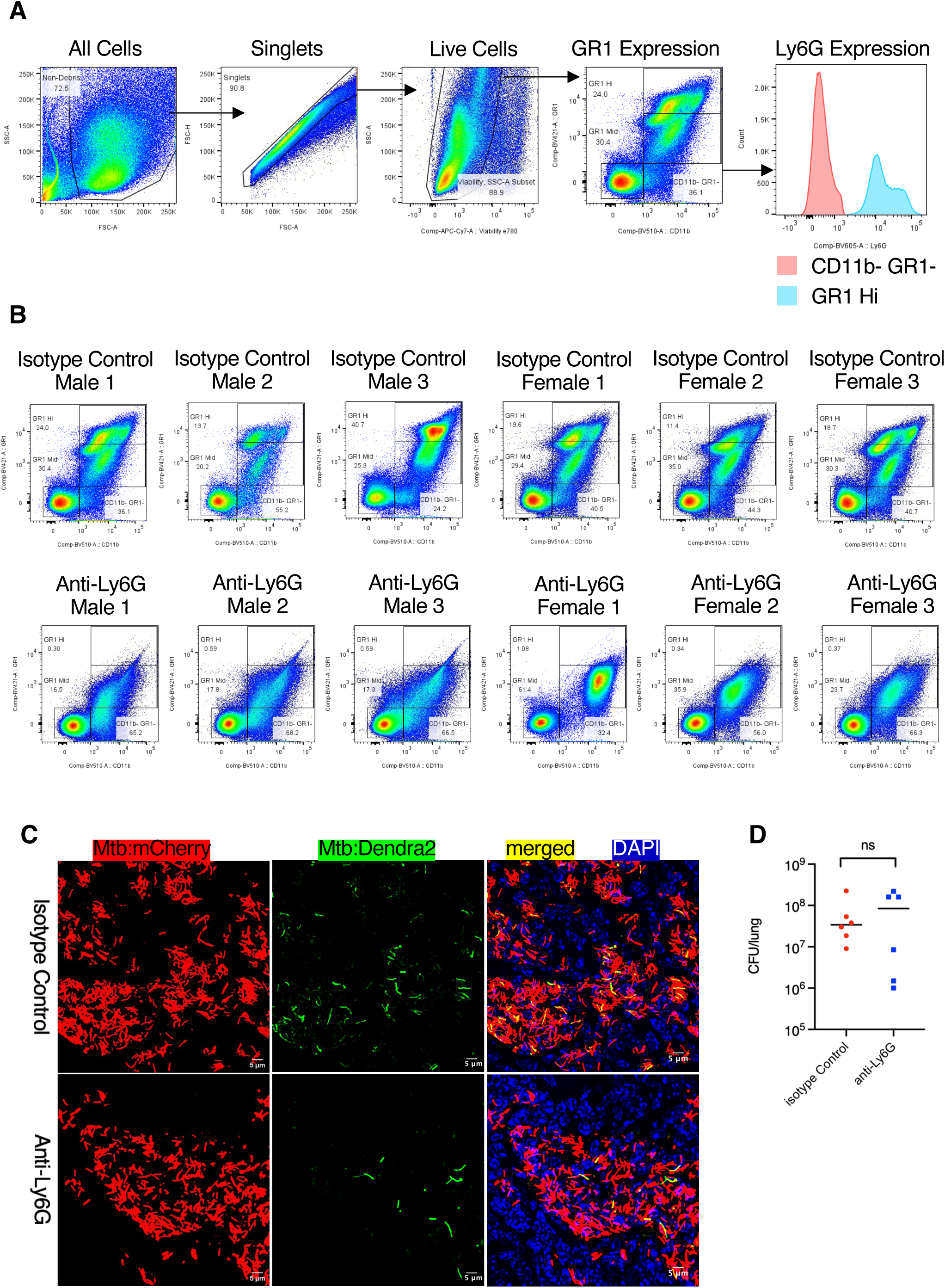
The effect of neutrophil depletion on zinc starvation response in Mtb in mouse lungs summarized in Figure 5. **A.** Gating strategy for quantifying neutrophil population in lung cells by flow cytometry after staining with the viability dye (Efluor 780), anti-CD11b (BV510), anti-GR1 (Pac Blue) and anti-Ly6G (BV605) antibodies. Shown is a representative sample of lung cells from a mouse infected for one week with Mtb(Erd):R1:R3 reporter strain and treated daily for two weeks with isotype control antibody. Subpopulation of CD11b positive cells with high levels of GR1 marker (GR1 Hi) were gated as neutrophils upon verification of Ly6G expression. The GR1 Hi gate was then applied to all samples from both control and anti-Ly6G groups of mice. **B.** Frequency of GR1 Hi (neutrophils) population in the indicated samples, referring to summarized data in figure 5F. **C.** Representative micrographs of lung sections from the above-described mouse groups showing Dendra2 expression from C- r-protein promoter in Mtb cells constitutively expressing mCherry. The frequency of these bacilli is summarized in figure 4E. **D.** Total bacterial burden in the lung of each mouse after treatment with anti-Ly6G or isotype control antibodies. ns denotes statistically not significant (Mann-Whitney)

**Table S1:**
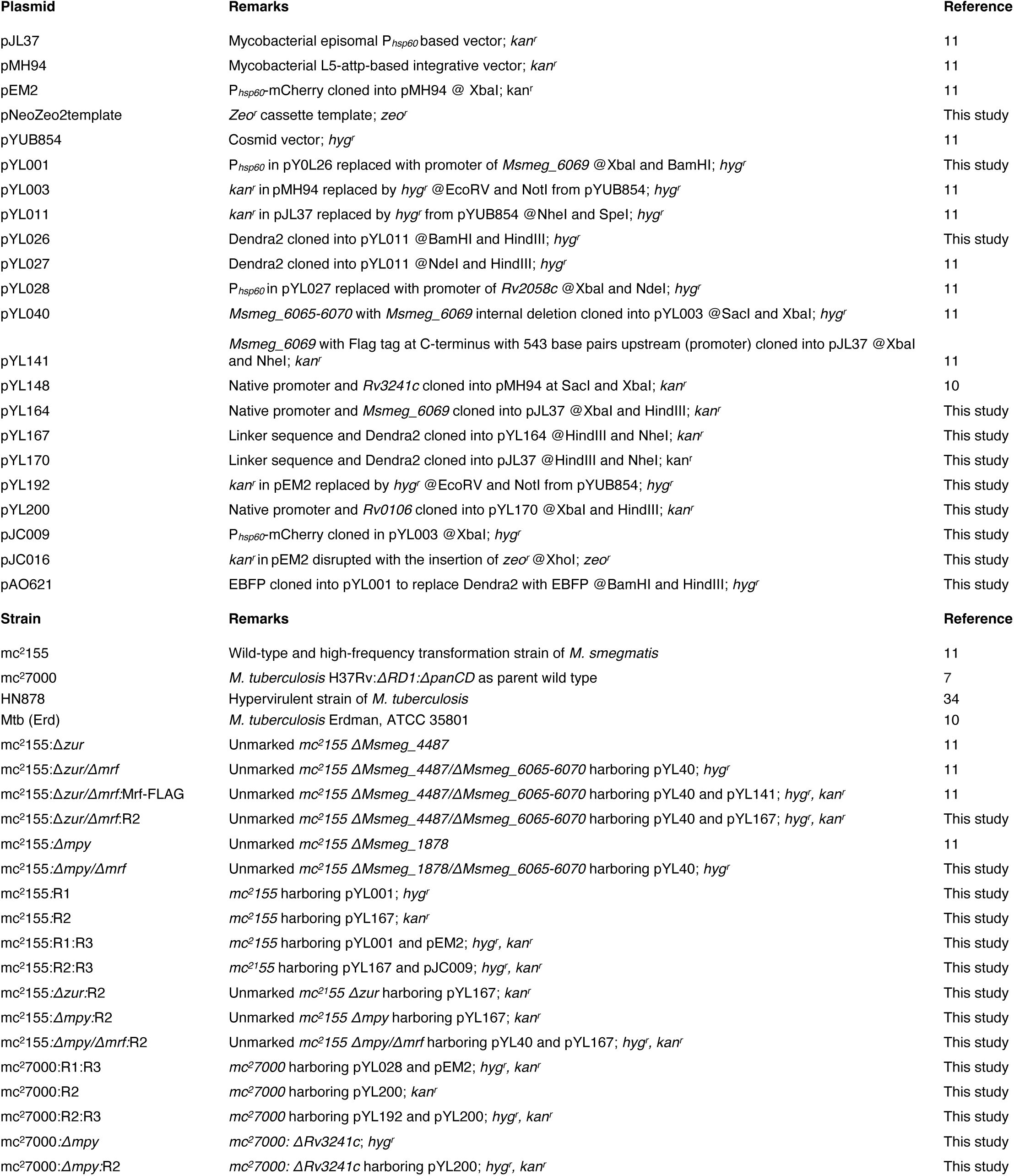
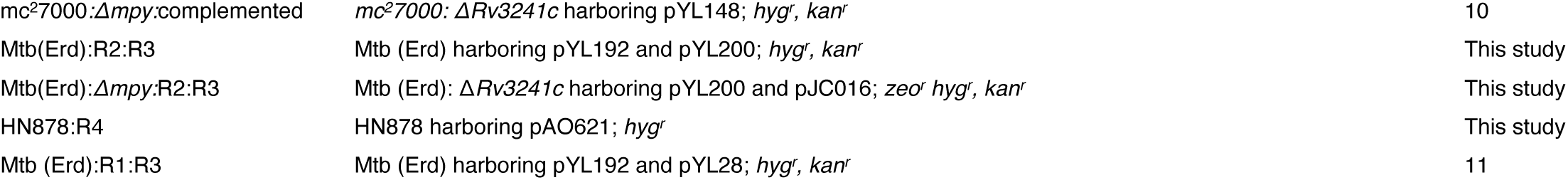
List of plasmids and strains used in this study.

**Table S2:**
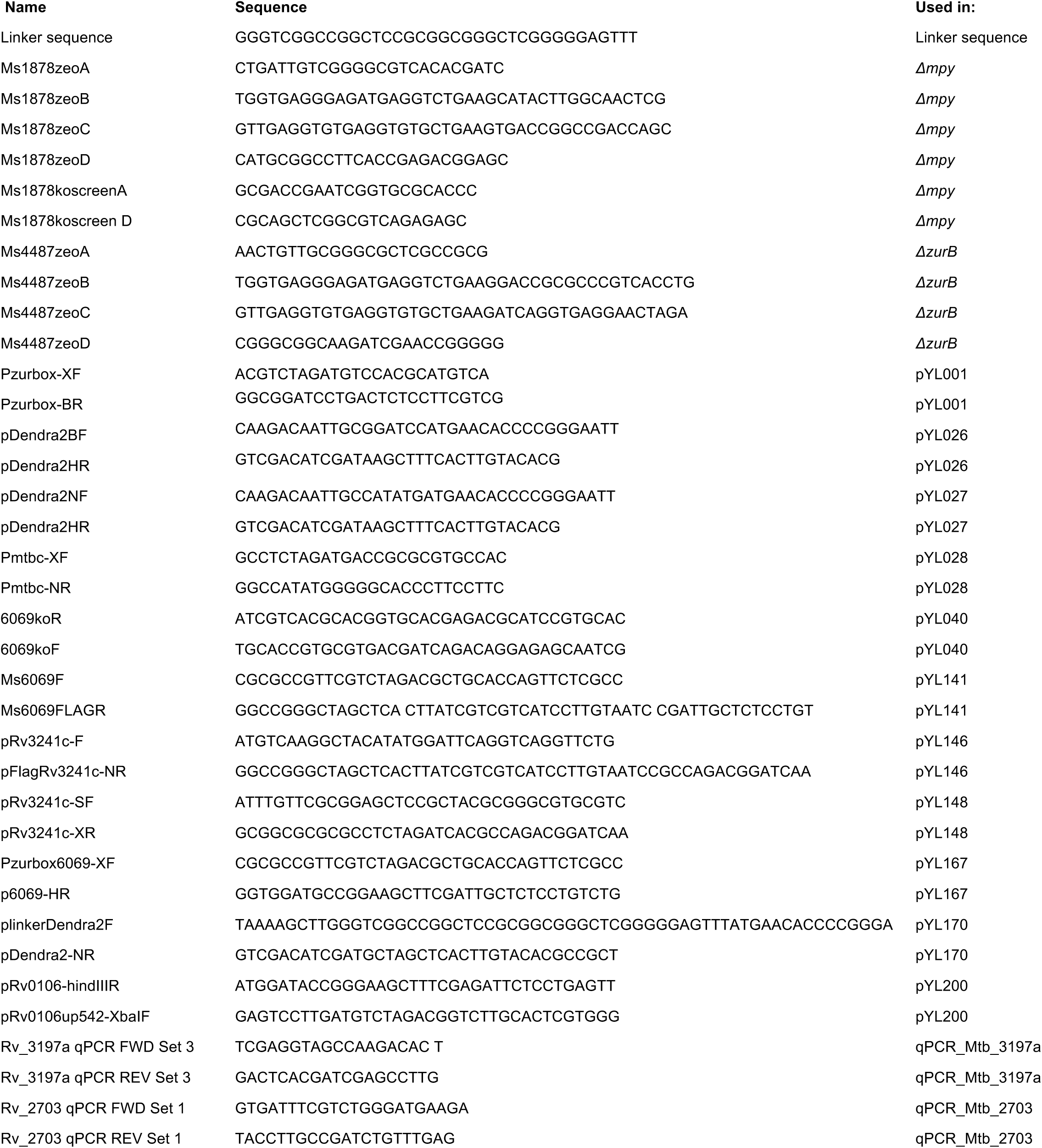
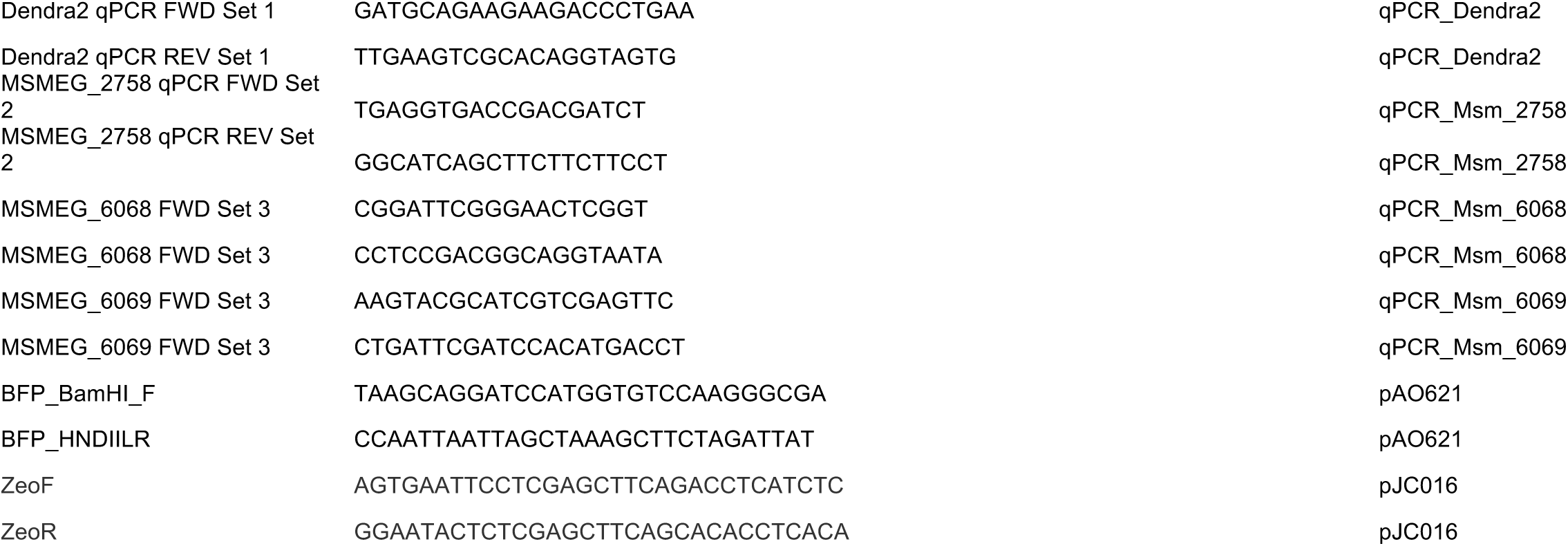
List of oligonucleotides.

## Notes

### Competing Interest Statement

The authors have declared no competing interest.

